# Population Genomics of the Bovine Sialic Acid Network Reveals Functional and Evolutionary Diversity Across European and South African Cattle Breeds

**DOI:** 10.64898/2026.11.06.764915

**Authors:** Oluwamayowa J. Ogun, Leke V. Aiyesa, Peter A. Idowu, Doreen Becker, Sebastian P. Galuska

## Abstract

**Background:** Sialic acids, mainly N-acetylneuraminic acid (Neu5Ac) and N-glycolylneuraminic acid (Neu5Gc), are terminal monosaccharides of glycans involved in immune and neuronal function. In bovine milk, they occur in sialylated oligosaccharides and other glycoconjugates, and their abundance varies among animals and changes rapidly during early lactation. Previous genetic studies have focused on single genes or individual milk traits, leaving variation across the genes involved in sialic acid biosynthesis, transport, transfer and turnover largely uncharacterized. We therefore characterized variation in 13 selected genes of the bovine sialic acid network using whole-genome sequence data from European and South African cattle.

**Results:** We analyzed 346 animals from 12 European and 4 South African breeds. Of 9,724 variants in the 13 target genes, 8,675 were intronic and 62 were protein-altering. Integration of SIFT predictions, Rate4Site conservation and protein context, with population representation as supporting evidence, identified 15 prioritized candidates, including 7 high-priority candidates in NEU3, ST8SIA2, ST8SIA4, CMAH and ST3GAL4. The most differentiated candidates were not among the 15 prioritized variants. NEU3 Y136H showed the largest regional contrast in allele frequency (0.83 in South African and 0.34 in European cattle), whereas several prioritized candidates were represented by a single allele in the South African cohort. Genome-wide principal component analysis of 950,219 linkage-disequilibrium-pruned SNPs separated South African from European cattle along the first component, which explained about 25% of the variance. None of the 220 pathway SNPs exceeded the genome-wide differentiation thresholds at breed, country or regional level. Within the ascertained pathway SNPs, diversity was higher in South African breeds, and Tajima’s D was shifted toward positive values in European breeds and centered near zero in South African breeds.

**Conclusions:** This study provides a network-wide map of protein-altering variation in the bovine sialic acid pathway and identifies 15 candidates for functional investigation, most of them rare. Because the most differentiated variants were not those with the strongest functional support, population contrasts, evolutionary conservation and protein context provide complementary evidence for selecting candidates. At the pathway level, differentiation remained within the genome-wide range. The candidates provide specific targets for genotyping, enzyme-level testing and genotype-informed milk phenotyping.

## 1. Introduction

Sialic acids are terminal monosaccharides that decorate soluble glycans, such as milk oligosaccharides (MOs), and glycan chains of RNA (Flynn et al., 2021), glycoproteins and glycolipids across mammalian tissues, including bovine milk. They are essentially involved in numerous physiological processes in mammals including the development of the immune and neuronal system. The two predominant mammalian sialic acid forms, N-acetylneuraminic acid (Neu5Ac) and N-glycolylneuraminic acid (Neu5Gc) (Varki and Varki, 2007), are present in cattle milk at concentrations that vary across breeds (Crisà et al., 2022; Sharma et al., 2019) and change dynamically during the early lactation period (Martín et al., 2001). Polysialic acid (polySia), a homopolymeric chain of Neu5Ac residues synthesized by the polysialyltransferases ST8 alpha-N-acetyl-neuraminide alpha-2,8-sialyltransferase 2 (ST8SIA2) and ST8 alpha-N-acetyl-neuraminide alpha-2,8-sialyltransferase 4 (ST8SIA4), shows a marked longitudinal decline during the first days of lactation in both bovine (Hinterseher et al., 2023) and ovine milk (Isernhagen et al., 2026b). The collective profile of sialic acid-containing glycans in milk, the milk sialoglycome, represents an important component of the bioactive glycan fraction of bovine milk.

This variation is not merely descriptive: Neu5Ac and Neu5Gc have distinct, and in some respects opposing, relevance to human health, giving the milk sialoglycome nutritional significance beyond cattle biology. Neu5Ac supports inter alia neurodevelopment and cognitive function (Wang, 2012), and Neu5Ac-containing glycans, including 3′-sialyllactose and 6′-sialyllactose, are added to some infant formulas (Li et al., 2025; Lasekan et al., 2022). Neu5Gc, by contrast, cannot be synthesized by humans because of the inactivation of cytidine monophosphate-N-acetylneuraminic acid hydroxylase (CMAH); functional CMAH catalyzes the hydroxylation of cytidine monophosphate (CMP)-Neu5Ac to CMP-Neu5Gc (Chou et al., 1998). However, Neu5Gc can be acquired through consumption of red meat and other foods from species that retain a functional CMAH enzyme, including pigs and cattle (Ogun et al., 2022; Tangvoranuntakul et al., 2003), where it is recognized as a xenoantigen. Interactions between dietary Neu5Gc and anti-Neu5Gc antibodies promoted inflammation and carcinoma development in Neu5Gc-deficient mice (Samraj et al., 2015), providing a proposed mechanism linking dietary Neu5Gc to inflammatory and pro-tumorigenic processes (Dhar et al., 2019). Bovine milk is an additional dietary route of Neu5Gc exposure (Tangvoranuntakul et al., 2003; Samraj et al., 2015; Crisà et al., 2022), so the genetic variants that may influence the Neu5Ac/Neu5Gc balance in milk are not only a question of cattle glycobiology, but a potential determinant of dietary xenoantigen exposure in humans.

Experimental quantification of Neu5Gc and Neu5Ac across seven European cattle breeds demonstrated substantial breed-specific variation in skeletal muscle sialic acid composition (Contò et al., 2025), suggesting that natural genetic background may influence sialic acid metabolism, yet whether similar mechanisms shape the milk sialoglycome specifically remains largely uncharacterized. The scale of this uncertainty is illustrated by sheep, where the Neu5Ac/Neu5Gc ratio is inverted relative to cattle and other mammals such as water buffalo, with Neu5Gc constituting approximately 90% of milk sialic acid rather than being the minority form as in cattle (Isernhagen et al., 2026a). The question of which specific enzyme variants, regulatory differences, or population-level allele frequency patterns drive these breed- and species-level differences in the milk sialoglycome remains a fundamental gap that motivates the present study.

As the milk sialoglycome depends on the coordinated activity of the sialic acid biosynthesis pathway, understanding the genetic variation within this network represents a logical first step towards identifying the molecular basis of breed-specific differences. The sialic acid biosynthesis network in cattle comprises a coordinated sequence of enzymatic reactions converting uridine diphosphate N-acetylglucosamine (UDP-GlcNAc) through a series of intermediates to produce CMP-Neu5Ac, which serves as an activated sialic acid donor substrate for sialyltransferases in the Golgi apparatus (Schauer, 2000). The rate-limiting enzyme UDP-N-acetylglucosamine 2-epimerase/N-acetylmannosamine kinase (GNE) initiates the pathway, followed by N-acetylneuraminate synthase (NANS), N-acetylneuraminate-9-phosphatase (NANP), and cytidine monophosphate N-acetylneuraminic acid synthetase (CMAS), with solute carrier family 35 member A1 (SLC35A1) transporting the activated substrate into the Golgi compartment. As mentioned above, CMAH catalyzes the hydroxylation of CMP-Neu5Ac to CMP-Neu5Gc, thereby determining the Neu5Gc/Neu5Ac balance. Sialyltransferases including beta-galactoside alpha-2,6-sialyltransferase 1 (ST6GAL1), ST3 beta-galactoside alpha-2,3-sialyltransferase 1 (ST3GAL1), and ST3 beta-galactoside alpha-2,3-sialyltransferase 4 (ST3GAL4) transfer nucleotide-activated sialic acid residues to specific glycan positions on milk glycans. In addition, ST8SIA2 and ST8SIA4 use the CMP-Neu5Ac substrate pool to synthesize polysialic acid (Sato et al., 2016). Moreover, the sialidases neuraminidase 1 (NEU1) and neuraminidase 3 (NEU3) are able to cleave sialic acid residues. Natural genetic variation across any node in this network could alter flux, substrate availability, or product specificity, and thereby shape the final sialoglycome composition of milk, a principle well established in human metabolic pathway genetics: GNE variants that reduce UDP-GlcNAc 2-epimerase/ManNAc kinase activity cause hyposialylation-associated myopathy (Eisenberg et al., 2001), and the Alu-mediated inactivation of human CMAH eliminated endogenous Neu5Gc synthesis in humans (Chou et al., 1998).

Population genomics provides a powerful framework for characterizing the landscape of natural genetic variation across biological pathways without requiring experimental manipulation. Large-scale cattle genomic resources have substantially expanded the availability of whole-genome sequence data across cattle populations. More recently, publicly available whole-genome resources spanning European and African cattle populations have enabled breed-stratified analysis of natural genomic variation at high resolution (Ginja et al., 2025; Dlamini et al., 2026). South African cattle breeds, including Nguni, Afrikaner, Bonsmara, and Tuli, represent populations with distinct demographic, environmental, and breeding histories and carry genomic diversity that complements that represented in European breed panels. To our knowledge, no study has applied population genomics to the sialic acid biosynthesis network in any cattle population. Here we address this gap by characterizing protein-altering variation across the bovine sialic acid network in European and South African cattle breeds.

## 2. Materials and Methods

### 2.1 Target Gene Selection and Genomic Coordinates

Thirteen genes representing major functional components of the bovine sialic acid biosynthesis, transport, modification, transfer, polysialylation, and turnover network were selected as targets for variant discovery (Table 1). Genes were classified into four functional tiers: (i) core biosynthesis and activation pathway, GNE, NANS, NANP, CMAS, SLC35A1, and CMAH; (ii) polysialyltransferases utilizing CMP-Neu5Ac as donor substrate, ST8SIA2 and ST8SIA4; (iii) terminal sialyltransferases, ST6GAL1, ST3GAL1, and ST3GAL4; and (iv) sialidases involved in sialic acid turnover, NEU1 and NEU3 (Varki and Schauer, 2009; Wickramasinghe and Medrano, 2011; Ogun et al., 2022). ST6GAL1, ST3GAL1, and ST3GAL4 were included as representative α2,6- and α2,3-sialyltransferases based on the documented transcriptional activity of these sialyltransferase classes in bovine milk somatic cells, including expression of ST3GAL-family genes and ST6GAL1, together with evidence of transcript structure for ST3GAL1 and ST6GAL1 (Wickramasinghe et al., 2011). The selected genes therefore represent biologically relevant control points rather than an exhaustive survey of the bovine sialyltransferase repertoire.

**Table 1.** Target genes for variant discovery, their function in the sialic acid biosynthesis network, functional category, and genomic position in the ARS-UCD1.2 bovine reference genome assembly (Ensembl release 110).

| Gene | Function in Sialoglycome | Tier | Genomic Position |
|---|---|---|---|
| GNE | Rate-limiting step: UDP-GlcNAc → ManNAc | 1, Core | BTA8:g.60,494,277-60,528,253 (−) |
| NANS | ManNAc → Neu5Ac-9-P | 1, Core | BTA8:g.63,192,882-63,212,682 (+) |
| NANP | Neu5Ac-9-P → Neu5Ac | 1, Core | BTA13:g.10,697,268-10,705,509 (+ |
| CMAS | Neu5Ac → CMP-Neu5Ac | 1, Core | BTA5:g.88,154,132-88,170,546 (−) |
| SLC35A1 | CMP-Neu5Ac Golgi transporter | 1, Core | BTA9:g.62,413,961-62,445,501 (−) |
| CMAH | CMP-Neu5Ac → CMP-Neu5Gc | 1, Core | BTA23:g.32,672,258-32,753,209 (+) |
| ST8SIA2 | PolySia synthase, uses CMP-Neu5Ac as donor substrate | 2, PolySia | BTA21:g.14,523,634-14,595,969 (−) |
| ST8SIA4 | PolySia synthase, uses CMP-Neu5Ac as donor substrate | 2, PolySia | BTA7:g.100,037,116-100,137,609 (−) |
| ST6GAL1 | α2-6 sialylation of glycoproteins | 3, Terminal | BTA1:g.80,194,829-80,267,738 (−) |
| ST3GAL1 | α2-3 sialylation | 3, Terminal | BTA14:g.7,865,552-7,951,420 (+) |
| ST3GAL4 | α2-3 sialylation | 3, Terminal | BTA29:g.29,698,724-29,750,191 (+) |
| NEU1 | Lysosomal sialidase, sialic acid turnover | 4, Turnover | BTA23:g.27,494,530-27,498,491 (+) |
| NEU3 | Plasma membrane sialidase, sialic acid turnover | 4, Turnover | BTA15:g.54,252,856-54,267,975 (+) |

*Reference: ARS-UCD1.2 bovine reference genome assembly.*

Genomic coordinates for all 13 target genes were defined using the Bos taurus ARS-UCD1.2 reference genome assembly (GCF_002263795.1). Gene boundaries were obtained from Ensembl release 110 and extended by 2 kb upstream, according to strand orientation, to capture proximal regulatory variation. The resulting intervals were used consistently for variant extraction and downstream analyses.

This panel represents a hypothesis-driven set of major mechanistic control points in sialic acid biosynthesis, activation, transport, Neu5Gc formation, terminal sialylation, polysialylation, and sialic acid turnover. It was not intended as an exhaustive catalog of genes capable of influencing the bovine sialoglycome.

### 2.2 Genomic Datasets

#### 2.2.1 European Cattle

Whole-genome sequencing data for European cattle were obtained from a publicly available resource generated by the OPTIBOV consortium (Ginja et al., 2025), comprising 289 animals from 11 native breeds and the commercial Holstein-Friesian breed across Finland, the Netherlands, and Portugal. Genomic DNA was sequenced on the Illumina NovaSeq6000 platform (paired-end, 150 bp reads) to an average depth of approximately 10× and mapped to the ARS-UCD1.2 reference genome (mean alignment rate 99.7%) using BWA-MEM2. Population-level variant calling was performed with FreeBayes followed by joint genotyping, and variants were annotated with the Ensembl Variant Effect Predictor.

Of the 289 animals represented in the published metadata, 226 animals across all 12 breeds could be unambiguously reconciled with sample identifiers in the deposited joint-called VCF and were retained for breed-resolved analyses (Table 2). The reduction primarily affected the three Portuguese native breeds, for which a subset of metadata identifiers could not be unambiguously matched to VCF sample identifiers. Animals for which metadata-to-VCF identifier correspondence could not be established unambiguously were excluded from breed-resolved analyses.

**Table 2.** European cattle breeds represented in the analyzed dataset, drawn from the Ginja et al. (2025) resource.

| **Breed** | **Country** | **N (analyzed)** | **N (published)** |
|---|---|---|---|
| Northern Finncattle | Finland | 25 | 25 |
| Western Finncattle | Finland | 25 | 25 |
| Eastern Finncattle | Finland | 25 | 25 |
| Holstein-Friesian | Finland/Netherlands/Portugal | 15 | 15 |
| Deep Red | Netherlands | 24 | 24 |
| Dutch Belted | Netherlands | 23 | 23 |
| Dutch Friesian | Netherlands | 24 | 24 |
| Meuse-Rhine-Yssel | Netherlands | 23 | 23 |
| Groningen White Headed | Netherlands | 21 | 21 |
| Barrosã | Portugal | 15 | 28 |
| Mirandesa | Portugal | 4 | 27 |
| Mertolenga | Portugal | 2 | 29 |
| Total |  | 226 | 289 |

Raw sequencing reads are available in the European Nucleotide Archive under accession PRJEB90816, while the joint-called population variant dataset used in this study is deposited in the European Variation Archive under accession PRJEB98152.

#### 2.2.2 South African Cattle

Whole-genome sequencing data for South African cattle were obtained from a publicly available OPTIBOV consortium resource covering indigenous cattle from Egypt, Uganda, and South Africa (Dlamini et al., 2026). Sequencing and variant-calling procedures followed those of the European resource: Illumina NovaSeq6000 paired-end 150-bp sequencing at approximately 10× average depth, BWA-MEM2 alignment to ARS-UCD1.2 (mean mapping rate 99.2%), and FreeBayes population variant calling with joint genotyping.The present study used the complete South African component, comprising 120 animals from four breeds (Table 3); Egyptian (n = 25) and Ugandan (n = 95) animals from the wider resource were not included.

**Table 3.** South African cattle breeds represented in the analyzed dataset, drawn from the Dlamini et al. (2026) resource.

| **Breed** | **Country** | **N (analyzed)** |
|---|---|---|
| Afrikaner | South Africa | 40 |
| Nguni | South Africa | 40 |
| Bonsmara | South Africa | 20 |
| Tuli | South Africa | 20 |
| Total |  | 120 |

Raw sequencing data are available in the European Nucleotide Archive under project accession PRJEB90914. Variant call files are available in the European Variation Archive under project accessions PRJEB110435 (ARS-UCD1.2 primary dataset) and PRJEB94226 (ARS-UCD1.2 filtered dataset).

### 2.3 Variant Extraction and Quality Control

For each of the 13 target genes, the complete annotated gene interval (Table 1) was extended by 2 kb upstream according to strand orientation, and the corresponding region was extracted directly from the remote source VCF using BCFtools and the accompanying local index file. Extracted regions were then subset to the study samples for each cohort (226 European animals and 120 South African animals).

Extracted variants were normalized, after which biallelic SNPs were retained for quality control. Individual genotype calls with read depth below 4 (DP < 4) or genotype quality below 20 (GQ < 20) were set to missing rather than removing the affected site. Following genotype masking, sites with a call rate below 90% (missingness ≥10%) were excluded. No minor allele frequency filter was applied at this stage, thereby preserving breed-specific and low-frequency variants for downstream analyses.

Post-QC variant characterisation was performed separately for each cohort using VCFtools v0.1.16 (Danecek et al., 2011) to assess per-individual and per-site missingness, heterozygosity, and allele frequency, and BCFtools v1.9 (Danecek et al., 2021) to calculate the transition/transversion ratio.

### 2.4 Functional Annotation

#### 2.4.1 Variant Annotation and Consequence Classification

QC-passed variants were annotated using Ensembl Variant Effect Predictor (VEP) v110 (McLaren et al., 2016) in offline mode against the Bos taurus ARS-UCD1.2 reference assembly and Ensembl v110 cache. The corrected union dataset comprised 10,261 variants, of which 9,724 were assigned to the 13 target genes. A total of 58 VEP-native protein-altering variants were identified, and 4 additional CMAH variants were resolved using the corresponding RefSeq transcript, yielding 62 protein-altering candidates in total (58 missense, 2 missense- and-splice-region, 1 stop-gained, and 1 stop-lost) for downstream functional characterization.

The 4 CMAH variants falling outside the Ensembl gene model but within the corresponding RefSeq transcript (XM_024984024.1) were resolved by RefSeq-based codon translation. The annotation procedure was validated against previously characterized CMAH variants (Ogun et al., 2023a). These 4 RefSeq-resolved candidates lacked native VEP SIFT scores and were treated accordingly in the functional-impact analysis (Section 2.4.2).

#### 2.4.2 Functional Impact Prediction

Functional impact of the 60 missense-containing candidates was assessed using native Bos taurus SIFT (Ng & Henikoff, 2003) predictions generated with Ensembl v110 (McLaren et al., 2016). SIFT predictions were extracted for the same transcript used for consequence classification in Section 2.4.1, and low-confidence predictions were retained separately. The four RefSeq-resolved CMAH candidates lacked corresponding Ensembl transcript annotations and were therefore classified as SIFT-unavailable. The stop-gained and stop-lost candidates were not evaluated by SIFT.

#### 2.4.3 Evolutionary Conservation Analysis

Evolutionary conservation at candidate residues was assessed using a fixed panel of 18 phylogenetically distributed vertebrate species selected independently of candidate-site results. One-to-one orthologues were retrieved from Ensembl Compara release 116 (Herrero et al., 2016) for each target gene, while ambiguous one-to-many relationships were excluded. Comparative sequences were combined with the corresponding Ensembl release 110 bovine reference protein and aligned using MAFFT v7.526 (Katoh & Standley, 2013), thereby preserving the protein-coordinate system used for candidate annotation. Candidate residues were considered assessable when at least six comparative orthologues contained a non-gap residue at the corresponding alignment position (Ashkenazy et al., 2016). The four RefSeq-resolved CMAH candidates were analyzed separately using the 577-aa RefSeq isoform X2 (XP_024839792.1) as the bovine reference.

Residue-specific evolutionary rates were estimated using Rate4Site v3.0.0 (Pupko et al., 2002) under the JTT substitution model with empirical Bayesian inference and a Gamma prior with 16 discrete categories. Lower Rate4Site scores indicate slower evolutionary rates and stronger relative conservation.

#### 2.4.4 Protein and Functional Context Annotation

Reference proteins used for candidate-site annotation were reconciled against bovine UniProtKB (UniProt Consortium, 2023) using sequence-identity testing and, where whole- sequence identity was not obtained, pairwise MAFFT alignment (Katoh & Standley, 2013). Direct residue-coordinate transfer from an external record was permitted only where exact sequence identity or explicit alignment-based reconciliation was established. UniProtKB features and InterPro domain/family annotations (Paysan-Lafosse et al., 2023) were evaluated for direct overlap with candidate residues. Overlapping annotations from multiple databases were not treated as independent evidence or combined into a numerical score. For ST3GAL4, seven bovine UniProtKB representations were independently reconciled because no single record provided a defensible equivalent of the candidate-defining reference protein. The four RefSeq-resolved CMAH candidates were evaluated separately by InterProScan v5.78-109.0 using the exact XP_024839792.1 sequence.

Candidates without reliable residue-level mapping were retained with protein-level context only. Absence of an annotated feature was not interpreted as evidence of functional irrelevance, and no structural modelling, docking, or molecular dynamics simulation was performed.

### 2.5 Population- and Breed-Level Allele Frequency Analysis

Allele frequencies for the 62 protein-altering candidates were calculated across the South African and European cohorts and separately for the 16 constituent breeds. Candidate identity was verified against each VCF record before extraction. Frequencies were derived from callable diploid genotypes after applying the established genotype-level thresholds (DP ≥ 4 and GQ ≥ 20), with missing genotypes excluded. Sites that failed the standard site-level call-rate threshold were still assigned a frequency from the genotype-masked callable subset, explicitly flagged as sub-threshold; sites entirely absent from the source variant dataset were not assigned an allele frequency, since homozygous-reference status could not be established from variant-only VCFs.

Breed-level allele counts were reconciled with the corresponding cohort-level estimates. Estimates based on fewer than five callable animals were flagged as very small-sample observations and retained for completeness but were not used as strong evidence of breed-specific enrichment

### 2.6 Integrated Candidate Prioritization

Evidence from functional prediction, evolutionary conservation, protein/domain context, pathway relevance, and population- and breed-level frequencies was integrated across all 62 candidates using chromosome, position, REF, and ALT alleles as the canonical variant identity. No arithmetic composite score was applied; evidence dimensions were retained separately, and candidates were prioritized based on convergence across complementary evidence types. Molecular consequence and functional prediction, conservation, and residue/domain context formed the primary functional evidence, while population and breed segregation was used as supporting evidence for downstream genotype-phenotype testing. Missing or non-assessable evidence was not treated as negative, and discordant evidence was retained rather than resolved by averaging.

### 2.7 Genome-Wide Population-Genomic Analyses

#### 2.7.1 Genome-Wide Dataset Preparation and Quality Control

Whole-genome variant datasets from the South African (Dlamini et al., 2026) and European (Ginja et al., 2025) cohorts described above, comprising 346 animals (120 South African and 226 European), were used for genome-wide population-genomic analyses. Biallelic SNPs were retained from each VCF file, and the two files were then merged, yielding 44,932,890 SNPs. Sites with more than 1% missing genotype calls were excluded, corresponding to a minimum site call rate of 99%, which reduced the dataset to 19,408,954 SNPs. A minor allele frequency (MAF) threshold of 0.05 was subsequently applied, leaving 12,954,798 SNPs. Genotype-level quality filtering required a minimum sequencing depth of 4 (DP ≥ 4) and genotype quality of at least 20 (GQ ≥ 20); following this step, the number of retained SNP loci remained 12,954,798. Variant filtering was performed using BCFtools (Danecek et al., 2021).

To reduce redundancy among correlated markers, linkage disequilibrium (LD) pruning was performed using PLINK v1.9 (Chang et al., 2015), with a step size of five variants and an r^2^ threshold of 0.25. The resulting LD-pruned dataset contained 950,219 SNPs on the 29 autosomes and formed the genome-wide marker set for subsequent population-genomic analyses.

Analyses of the 13 sialic-acid pathway genes used the candidate-gene datasets of Section 2.3 rather than the genome-wide dataset. After per-gene quality control in each cohort, the European and South African datasets were intersected with BCFtools (Danecek et al., 2021), and the 4,505 biallelic SNP records present in both cohorts with identical alleles were merged into a single gene-level dataset for the 346 animals. These records cover 4,504 unique positions, because one CMAH position (chr23:32,681,432) carries two alternative alleles recorded as separate biallelic SNPs. The 10,261 variants of the candidate-gene analysis represent the union of the two cohorts and therefore also include variants observed in only one cohort.

#### 2.7.2 Population Structure

Genetic structure among the 346 cohort animals was investigated by principal component analysis (PCA) using PLINK (Chang et al., 2015), based on the 950,219 LD-pruned SNPs. Principal components were examined to characterize genetic structure and differentiation between South African and European cattle and among individual breeds, and the resulting clustering patterns were compared with breed and country assignments to assess the extent to which genetic relationships corresponded to the predefined population groups.

#### 2.7.3 Genetic Diversity

Genetic diversity was characterized within breeds using the 4,505 SNP records of the gene-level dataset (Section 2.7.1) for the 346 animals of the 16 breeds. Minor allele frequency (MAF) was calculated from genotype counts for each SNP record and breed. Observed heterozygosity (H_O_) was calculated as the proportion of heterozygous genotypes among called genotypes, and expected heterozygosity (H_E_) as 2p(1 − p), where p is the alternate allele frequency among called diploid genotypes, without finite-sample correction. Nucleotide diversity (π) was calculated for each SNP record as π = [n/(n − 1)] × [1 − Σpᵢ²], where n is the number of called alleles and pᵢ is the frequency of allele i; records with fewer than two called alleles were excluded. Breed-level π was obtained as the mean across eligible SNP records. Because the dataset contains only SNP sites ascertained as variable in the candidate-gene analysis and was not filtered for MAF, these values represent mean diversity per ascertained SNP rather than nucleotide diversity per genomic base, and they describe the analyzed pathway regions only. Mertolenga (n = 2) and Mirandesa (n = 4) were retained in these descriptive summaries and flagged for cautious interpretation. All diversity measures were calculated with custom Python scripts.

#### 2.7.4 Population Differentiation

Population differentiation was assessed at breed, country, and regional levels using the Weir and Cockerham (1984) estimator implemented in VCFtools (Danecek et al., 2011). Population files were defined for individual breeds, countries (Finland, the Netherlands, Portugal, and South Africa), and the two regions (Europe and South Africa). Per-SNP F_ST_ was estimated jointly across all groups at each level on the 950,219 LD-pruned SNPs. Pathway SNPs were defined as LD-pruned SNPs located within the coordinates of the 13 genes, including the 2-kb upstream extension. Genome-wide outlier SNPs were defined at each level by the empirical 99.9th percentile (top 0.1%) of the per-SNP F_ST_ distribution (breed level 0.452, country level 0.475, regional level 0.580). Percentiles were calculated from finite estimates (949,858 at breed level and 949,880 at country and regional levels), excluding undefined values. Negative finite F_ST_ estimates were retained for percentile calculations but displayed as zero in Figure 9. Pathway SNPs with F_ST_ ≥ 0.25 were classified as highly differentiated for descriptive purposes. F_ST_ was interpreted descriptively, as a measure of differentiation between predefined groups, and not as a test of selection. For the 62 protein-altering candidate variants, genetic differentiation was additionally evaluated at regional, country and pairwise breed levels using the Weir and Cockerham (1984) F_ST_ estimator implemented in VCFtools, with genotypes extracted from the quality-controlled gene-specific VCF datasets. These candidate-level analyses were conducted separately from the genome-wide F_ST_ scan, and finite estimates were independently validated.

#### 2.7.5 Tajima’s D

Differences in the allele-frequency spectrum within the 13 sialic-acid pathway genes were evaluated using Tajima’s D (Tajima, 1989), implemented in VCFtools (Danecek et al., 2011). Tajima’s D compares two estimates of nucleotide diversity, one based on the average number of pairwise differences and one on the number of segregating sites, and measures departure from the neutral site-frequency spectrum. For each gene, the SNP records of that gene in the gene-level dataset (4,505 records in total; Section 2.7.1) were used as input, and only individuals belonging to the population or breed under investigation were retained. Tajima’s D was calculated in non-overlapping 1-Mb windows; because each gene lies within a single window, one value was obtained per gene and breed. The breed-level calculation included 22 Dutch Belted animals, one fewer than in the analyzed cohort (Table 2). Because the gene-specific datasets comprised segregating SNPs rather than complete resequencing alignments including invariant sites, Tajima’s D values were interpreted as comparative measures of the allele-frequency spectrum among populations rather than as direct tests of neutrality. Mertolenga (n = 2) and Mirandesa (n = 4) were excluded from the regional comparison because such small samples do not support reliable estimation of allele-frequency spectra.

### 2.8 Integration of Pathway Loci with Genome-Wide Evidence

Genome-wide population structure and differentiation were used as the demographic background against which variation in the 13 sialic-acid pathway genes was interpreted. Differentiation of pathway SNPs was compared with the genome-wide F_ST_ distribution at each level, whereas diversity and the allele-frequency spectrum were described within the pathway genes only, because no genome-wide estimates of these statistics were retained. Genes containing highly differentiated SNPs were compared with the genes carrying the prioritized candidates (Section 3.8). The gene-level dataset comprised 4,505 SNP records across the 13 sialic-acid pathway genes (Section 2.7.1). Candidate-level F_ST_ estimates (Section 2.7.4) were examined for the 62 protein-altering candidates to assess whether differentiation and functional prioritization coincided at individual variants. Candidates absent from the variant-only VCF records of a group were not imputed as reference genotypes, so F_ST_ was not estimable for those comparisons; finite and non-estimable comparisons are reported separately for each level, and raw negative estimates were retained. No formal haplotype-based test of selection, such as iHS or XP-EHH, was performed in this study, and differentiation and frequency-spectrum statistics are reported descriptively.

## 3 Results

### 3.1 Variant Landscape Across the Bovine Sialic Acid Biosynthesis Network

Following quality control and correction of the *CMAH* extraction boundary, the combined South African and European datasets contained 10,261 unique variants across the analyzed genomic intervals. Of these, 9,724 variants were assigned to the 13 target genes, while 537 annotations associated with neighboring non-target loci were excluded. The target-gene variants were predominantly intronic (n = 8,675), with additional variants occurring in upstream, downstream, untranslated, synonymous, splice-associated, and protein-altering regions.

Variant burden differed substantially among genes (Figure 1; Table 4). ST8SIA2 contained the largest number of target-gene variants (n = 1,632), followed by CMAH (n = 1,584), ST3GAL1 (n = 1,566), and ST8SIA4 (n = 1,435), whereas NEU1 contained the fewest (n = 84). After accounting for the length of the analyzed interval, including the 2-kb upstream extension, variant density was highest for ST8SIA2 (21.95 variants/kb), followed by CMAH (19.10 variants/kb) and ST3GAL1 (17.82 variants/kb), and lowest for SLC35A1 (8.86 variants/kb).

**Figure 1.**
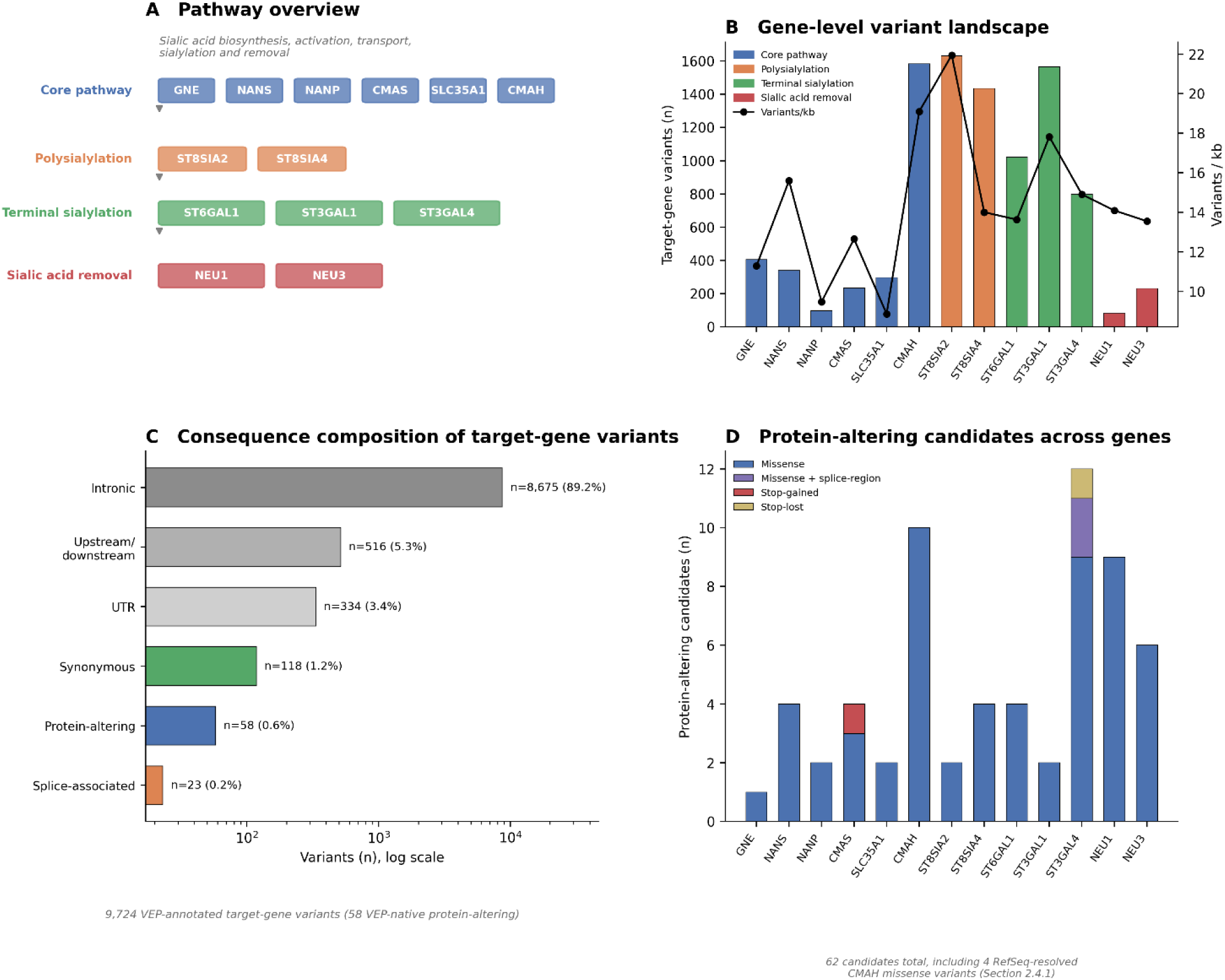
Variant landscape across the 13-gene bovine sialic-acid network. (A) Pathway overview across four functional tiers. (B) Target-gene variant counts and density per gene. (C) Consequence composition of the 9,724 VEP-annotated target-gene variants (58 VEP-native protein-altering). (D) Distribution of the final 62 protein-altering candidates, including four RefSeq-resolved CMAH variants (Section 2.4.1).

**Table 4.** Gene-level variant landscape.

| Gene | Functional tier | Analyzed interval (kb) | Target-gene variants | Variants/kb | Protein-altering candidates |
|---|---|---|---|---|---|
| GNE | Core | 35.977 | 406 | 11.29 | 1 |
| NANS | Core | 21.801 | 340 | 15.60 | 4 |
| NANP | Core | 10.242 | 97 | 9.47 | 2 |
| CMAS | Core | 18.415 | 233 | 12.65 | 4 |
| SLC35A1 | Core | 33.541 | 297 | 8.86 | 2 |
| CMAH | Core | 82.952 | 1,584 | 19.10 | 10 |
| ST8SIA2 | PolySia | 74.336 | 1,632 | 21.95 | 2 |
| ST8SIA4 | PolySia | 102.494 | 1,435 | 14.00 | 4 |
| ST6GAL1 | Terminal | 74.910 | 1,021 | 13.63 | 4 |
| ST3GAL1 | Terminal | 87.869 | 1,566 | 17.82 | 2 |
| ST3GAL4 | Terminal | 53.468 | 797 | 14.91 | 12 |
| NEU1 | Recycling | 5.962 | 84 | 14.09 | 9 |
| NEU3 | Recycling | 17.120 | 232 | 13.55 | 6 |
| Total |  |  | **9,724** |  | **62** |

Protein-altering annotation identified 62 candidate variants across all 13 genes, comprising 58 missense variants, two missense variants overlapping splice regions, one stop-gained variant, and one stop-lost variant. Fifty-eight candidates were identified directly from the Ensembl-based annotation, while four additional CMAH missense variants were recovered through RefSeq-based annotation of the corrected CMAH interval. Candidate burden was highest in ST3GAL4 (n = 12), followed by CMAH (n = 10), NEU1 (n = 9), and NEU3 (n = 6), while all 13 pathway genes contained at least one protein-altering candidate.

### 3.2 Population and Breed-Level Distribution of Protein-Altering Variants

The 62 protein-altering candidates showed distinct patterns of representation and allele frequency between the South African and European cohorts. 31 candidates were retained as callable sites in both cohorts. A further 21 candidates were present in the South African cohort but were not represented in the European source variant dataset, whereas 10 candidates were present in Europe but were either not represented in the South African source dataset (n = 6) or did not pass the South African site-level quality-control threshold (n = 4). Because absence from a variant-only source VCF does not establish homozygous-reference status, variants not represented in the source dataset were not assigned an allele frequency.

Among the 31 candidates retained in both cohorts, allele frequencies varied substantially between populations (Figure 2A). The largest difference occurred for the NEU3 variant at chr15:54265634, with an allele frequency of 0.8276 in South African cattle compared with 0.3447 in European cattle (absolute ΔAF = 0.4829). Smaller but notable differences were observed for 3 ST3GAL4 candidates at chr29:29730119, chr29:29730201, and chr29:29730070 (absolute ΔAF = 0.1230, 0.1104, and 0.1089, respectively), and for a NANS candidate at chr8:63193083 (absolute ΔAF = 0.1003).

**Figure 2.**
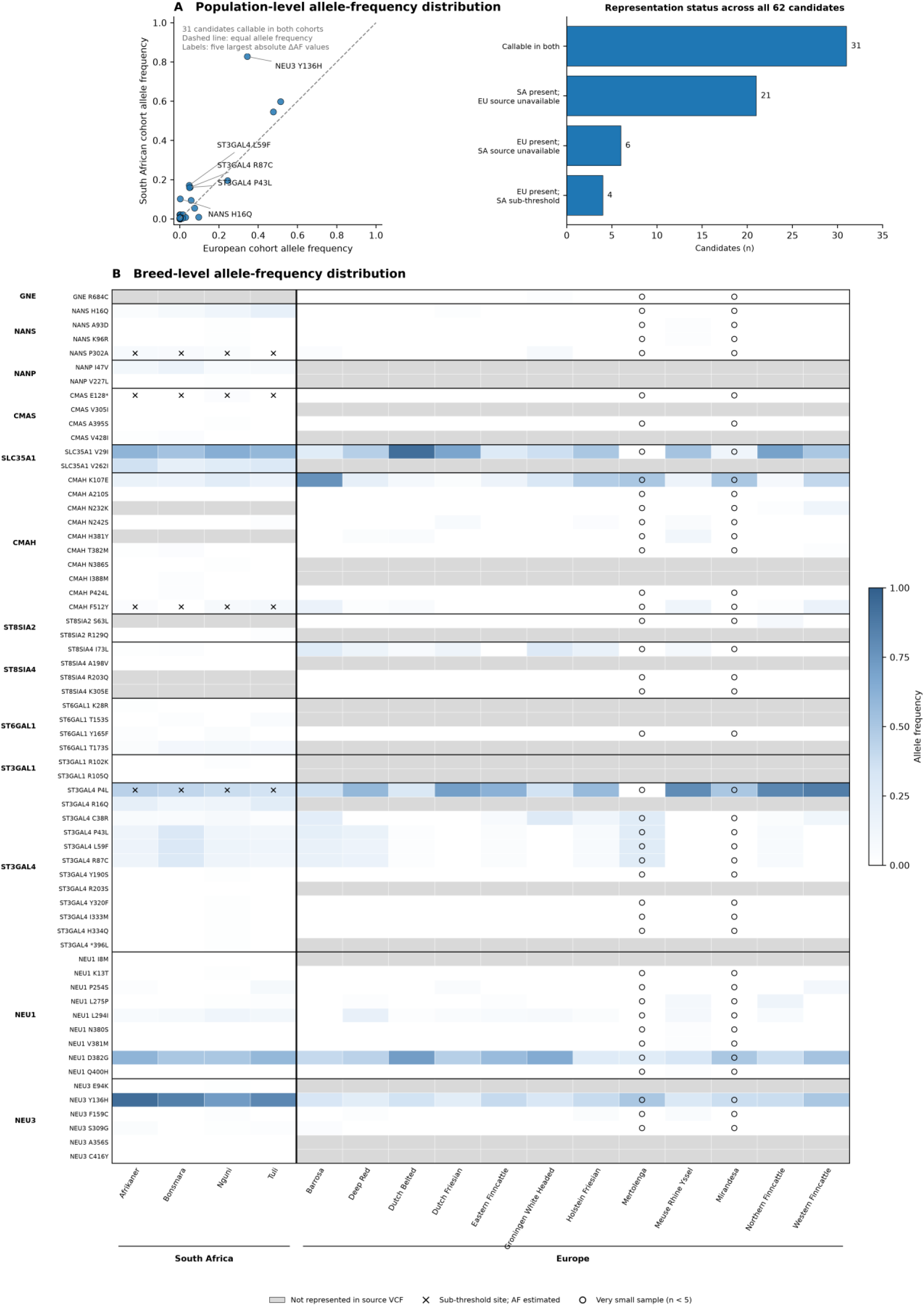
Population- and breed-level allele-frequency distribution of the 62 protein-altering candidates. (A) South African versus European cohort allele frequency for the 31 candidates callable in both cohorts (left), with a compact summary of representation status across all 62 candidates (right); the dashed line indicates equal frequency between cohorts, and the five candidates with the largest absolute allele-frequency differences are labelled. (B) Breed-level allele frequency for all 62 candidates across the 4 South African and 12 European constituent breeds. Grey cells indicate variants not represented in the source VCF for that breed; × marks sub-threshold sites retained from callable genotypes after the site-level quality-control threshold was not met; ○ marks estimates from very small samples (Mertolenga, n = 2; Mirandesa, n = 4). The complete numerical matrix underlying this panel is provided in Supplementary Table S1.

Breed-level analysis further revealed heterogeneous distributions among the 16 constituent breeds (Figure 2B; Supplementary Table S1). Across the 992 candidate-breed combinations, 700 were classified as present, 16 were absent after application of the standard site-level quality-control threshold but retained as sub-threshold frequency estimates from callable genotypes, and 276 were not represented in the corresponding source variant dataset. Across the 716 combinations with estimable frequencies, 634 involved adequately sampled breeds (n ≥ 5 callable animals), whereas 82 involved very small samples, all confined to Mertolenga (n = 2) and Mirandesa (n = 4); these estimates are retained for completeness but are not used as strong evidence of breed-specific enrichment.

Among reliably sampled breed-level estimates, 14 candidates were observed in all 4 South African breeds, whereas 19 were observed in only 1 South African breed. In the European cohort, 5 candidates were observed across all 10 breeds with standard sample-size support, while 10 were observed in only 1 such breed. Several broadly distributed candidates nevertheless showed marked frequency variation among breeds: CMAH chr23:32721570 ranged from 0.0417 to 0.7692 across European breeds, SLC35A1 chr9:62441325 from 0.2500 to 0.9286, and ST3GAL4 chr29:29729953 from 0.3043 to 0.8600. The strongly differentiated NEU3 chr15:54265634 variant was observed across all 4 South African breeds at frequencies of 0.7250-0.9324, compared with 0.2391-0.5000 across 10 European breeds. These population- and breed-level frequency patterns were treated as evidence of segregation rather than functional importance in themselves and were subsequently evaluated together with functional-impact prediction, evolutionary conservation, and protein-context evidence (Sections 3.3-3.6).

### 3.3 Predicted Functional Impact of Candidate Variants

Native bovine SIFT predictions were available through VEP for 56 of the 62 protein-altering candidates. The four RefSeq-resolved CMAH missense variants lacked native VEP SIFT annotations, while the CMAS E128* stop-gained and ST3GAL4 *396L stop-lost variants were not applicable to SIFT analysis. Among the 56 assessable variants, 6 were classified as deleterious, 6 as deleterious with low confidence, 38 as tolerated, and 6 as tolerated with low confidence (Figure 3). The 6 deleterious predictions comprised NEU3 E94K and F159C, ST8SIA2 R129Q, NEU1 L275P, CMAH N242S, and ST3GAL4 R203S, with SIFT scores ranging from 0.00 to 0.03. A further 6 candidates were classified as deleterious with low confidence: NEU1 I8M, ST3GAL4 P4L, R16Q, C38R, and P43L, and GNE R684C. Notably, 4 of the 6 low-confidence deleterious predictions occurred in ST3GAL4, indicating a local concentration of predicted functional sensitivity within this gene, although these classifications were retained separately from standard-confidence deleterious predictions.

**Figure 3.**
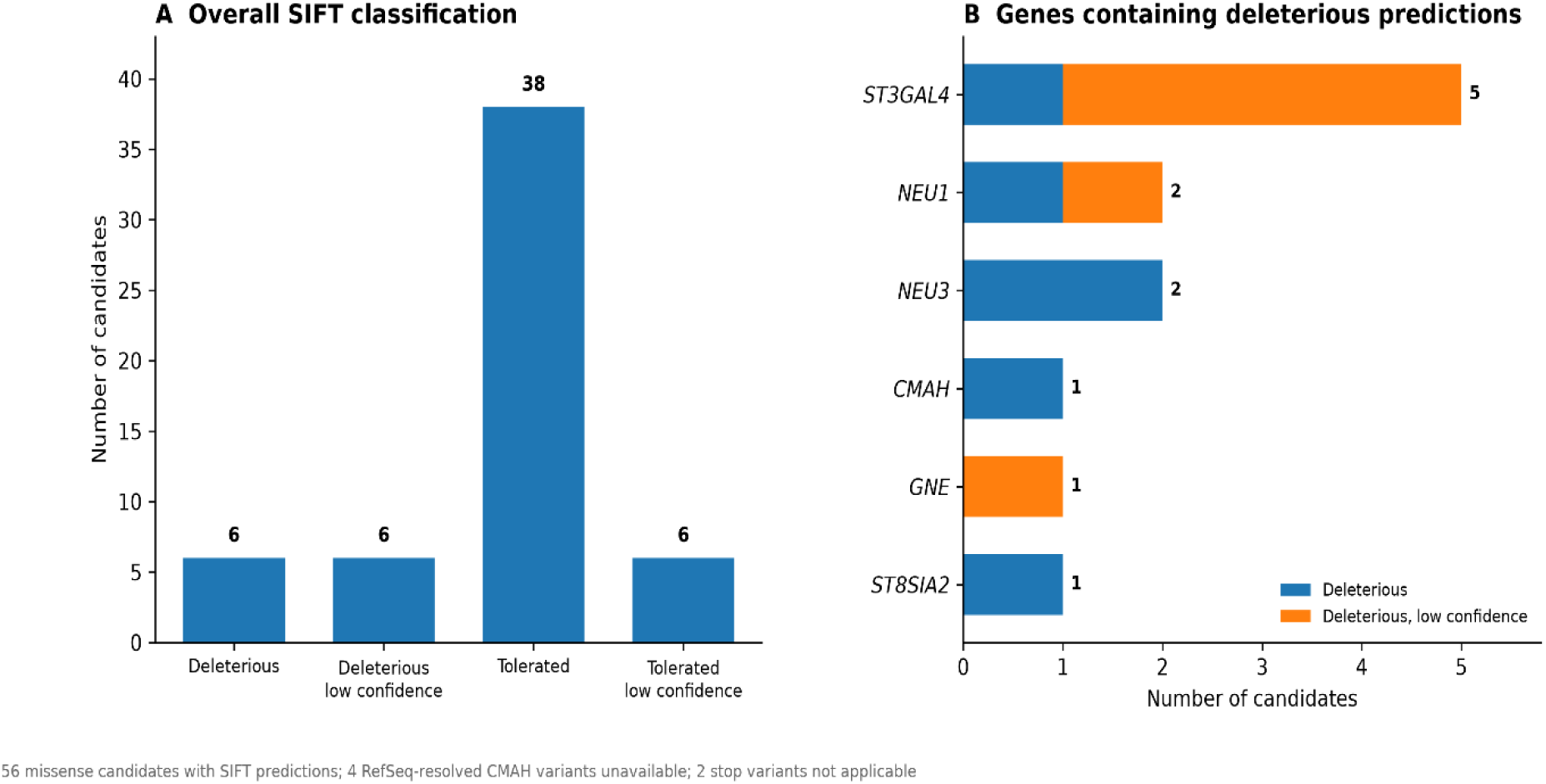
Predicted functional impact of the 62 protein-altering candidates based on SIFT. (A) Distribution of SIFT classifications among the 56 missense-containing candidates with native bovine SIFT predictions. Four RefSeq-resolved CMAH missense variants lacked native VEP SIFT annotations, and the CMAS E128* stop-gained and ST3GAL4 *396L stop-lost variants were not applicable to SIFT analysis. (B) Gene-level distribution of the 12 candidates classified as deleterious or deleterious with low confidence. Standard- and low-confidence deleterious predictions are shown separately.

Overall, most SIFT-assessable candidates were predicted to be tolerated, whereas a smaller subset showed evidence consistent with possible amino-acid-level functional disruption. Because SIFT predictions reflect sequence-based functional constraint rather than direct experimental effect, these results were subsequently evaluated together with evolutionary conservation and protein-context evidence in Sections 3.4–3.6.

### 3.4 Evolutionary Conservation of Candidate Residues

Evolutionary conservation analysis revealed substantial variation in residue-level constraint among the protein-altering candidates. Rate4Site scores were obtained for 57 of the 62 candidates, while 4 were not assessable and the ST3GAL4 396L stop-lost variant was treated as a separate special case. The strongest conservation signal was observed for ST8SIA4 I73L (Rate4Site = −0.9582), followed by ST8SIA2 R129Q (−0.9269), ST8SIA4 R203Q (−0.8719), CMAS E128* (−0.8396), NEU3 E94K (−0.6812), and the RefSeq-resolved CMAH I388M (−0.6392) (Figure 4).

**Figure 4.**
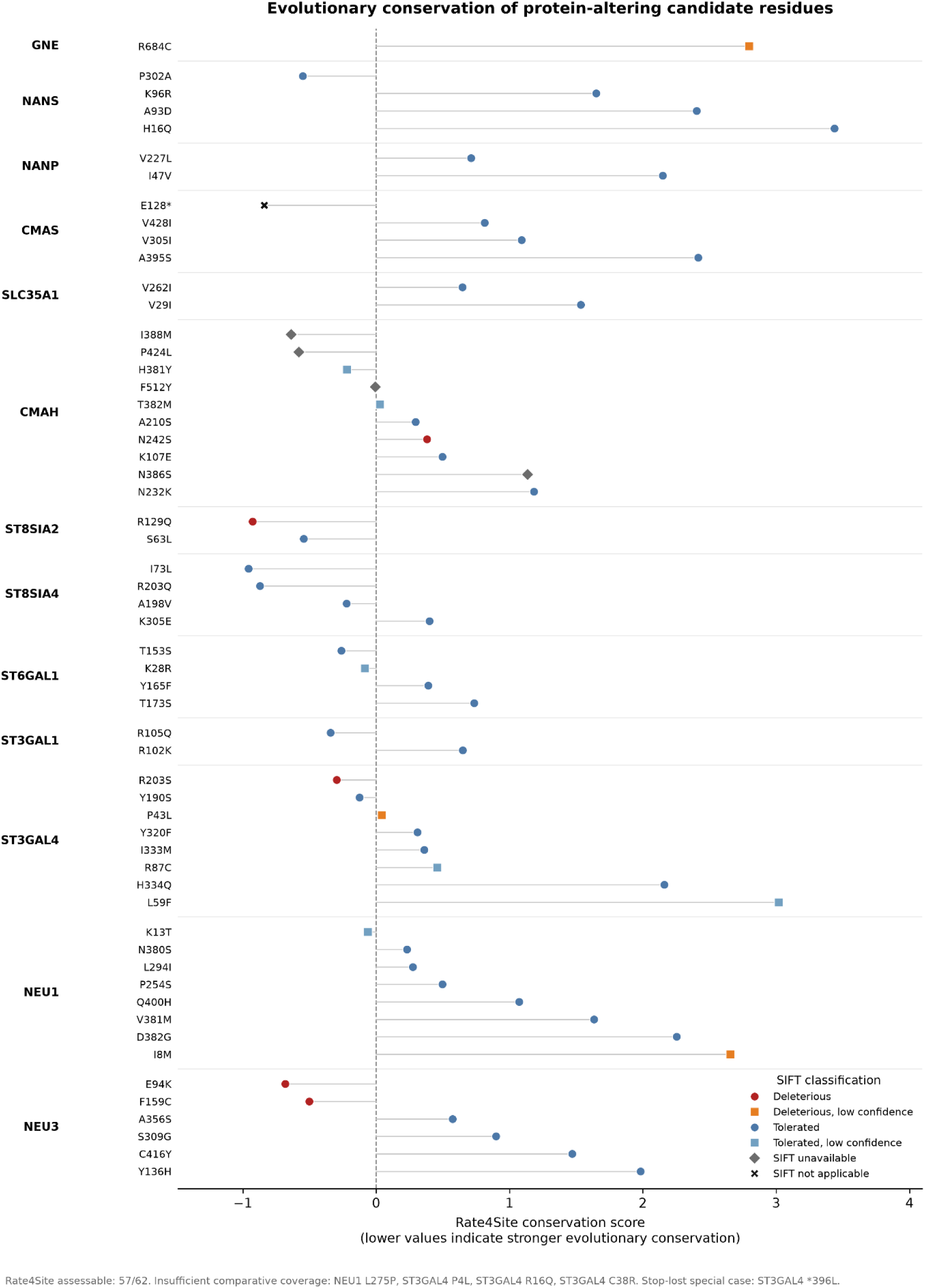
Evolutionary conservation of protein-altering candidate residues across the bovine sialic-acid network. Rate4Site conservation scores are shown for the 57 of 62 protein-altering candidates that met the predefined comparative-sequence coverage criterion. Candidates are grouped by gene, with lower Rate4Site scores indicating stronger evolutionary conservation. Marker color and shape indicate the corresponding SIFT classification, allowing conservation and predicted functional impact to be evaluated jointly. NEU1 L275P was not assessable because the candidate position fell within an alignment gap, whereas ST3GAL4 P4L, R16Q, and C38R had insufficient comparative sequence coverage. The ST3GAL4 *396L stop-lost variant was treated as a separate special case and was not assigned a Rate4Site score.

Several candidates showed concordance between evolutionary conservation and SIFT prediction. In particular, ST8SIA2 R129Q, NEU3 E94K and F159C, and ST3GAL4 R203S occurred at evolutionarily constrained residues and were also classified as deleterious by SIFT. Conversely, strongly conserved residues were also identified among candidates predicted as tolerated, including ST8SIA4 I73L and R203Q, illustrating the complementary information provided by the two approaches. The RefSeq-resolved CMAH I388M and P424L variants also occurred at relatively conserved residues despite lacking native VEP SIFT predictions. Four missense candidates could not be reliably evaluated for evolutionary conservation. NEU1 L275P could not be scored because the corresponding candidate position fell within an alignment gap, whereas ST3GAL4 P4L, R16Q, and C38R had insufficient comparative sequence coverage at the corresponding alignment positions to meet the predefined threshold. These candidates were retained rather than interpreted as unconserved. Overall, conservation evidence identified a subset of constrained candidate residues and provided an independent evidence dimension for subsequent protein-context analysis and integrated prioritization.

### 3.5 Protein and Functional Context of Candidate Residues

Protein and functional context was evaluated for all 62 candidates using UniProt and InterPro annotations (Figure 5; Supplementary Table S2). Residue-level context was obtained for 52 candidates: 43 through direct or alignment-reconciled mapping, 5 through multi-reference reconciliation of ST3GAL4, and 4 through sequence-based annotation of RefSeq-resolved CMAH variants. The remaining 10 candidates were retained at protein-level context only. Among the 43 straightforward mappings, 22 overlapped annotated UniProt features.

**Figure 5.**
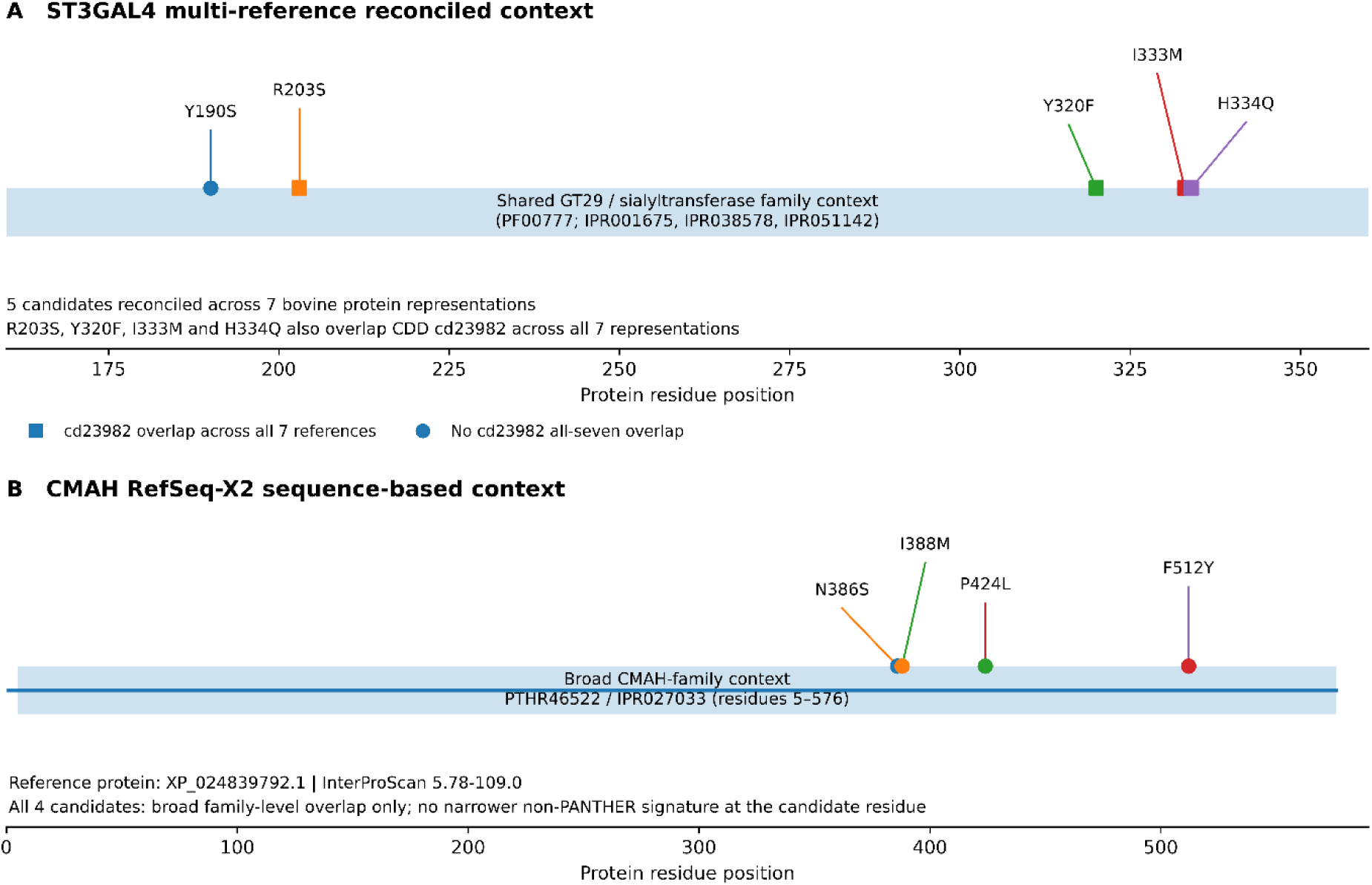
Protein and functional context of selected candidates requiring reference reconciliation. (A) Five ST3GAL4 candidates reconciled across seven bovine protein representations. All five overlapped shared GT29/sialyltransferase family annotations, while R203S, Y320F, I333M, and H334Q additionally overlapped CDD cd23982 across all seven representations. (B) Four RefSeq-X2 CMAH candidates mapped within the broad PTHR46522/IPR027033 family context on XP_024839792.1; no narrower non-PANTHER signature overlapped the candidate residues.

Five ST3GAL4 candidates (Y190S, R203S, Y320F, I333M, and H334Q) were reconciled across seven bovine protein representations and consistently overlapped broader glycosyltransferase/sialyltransferase annotations, including PF00777 and InterPro entries IPR001675, IPR038578, and IPR051142. Four (R203S, Y320F, I333M, and H334Q) additionally overlapped CDD cd23982 across all seven representations, whereas Y190S did not. Other ST3GAL4 candidates that could not be reliably reconciled at residue level were retained at protein-level context, with *396L treated separately as a stop-lost variant. The four RefSeq-resolved CMAH candidates (N386S, I388M, P424L, and F512Y) fell within the broad CMAH-family annotation (PTHR46522; IPR027033) but did not overlap a narrower non-PANTHER InterProScan signature at the candidate residue. They were therefore retained with broad family-level rather than specific domain-level context.

### 3.6 Integrated Prioritization of Candidate Variants

Integration of functional-impact prediction, evolutionary conservation, protein and domain context, pathway relevance, and population- and breed-level segregation identified 15 prioritized candidates from the 62 protein-altering variants (Figure 6; Supplementary Table S3). Seven were classified as high-priority and eight as secondary candidates based on qualitative convergence across complementary evidence dimensions. No arithmetic composite score was applied.

**Figure 6.**
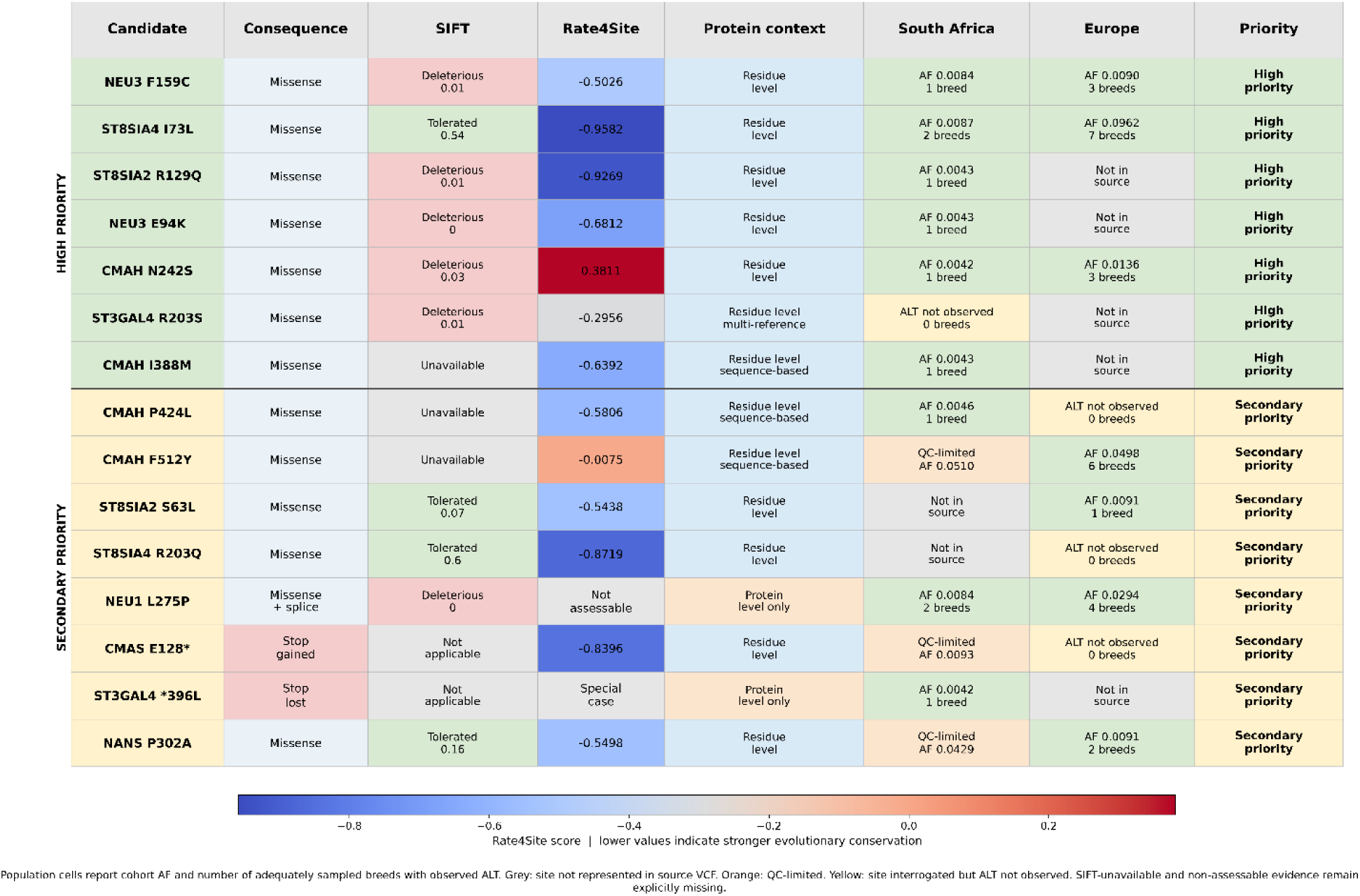
Integrated evidence supporting prioritization of 15 protein-altering candidates. Functional consequence, SIFT prediction, Rate4Site conservation, protein-context resolution, and South African and European population evidence are shown independently for seven high-priority and eight secondary-priority candidates. Lower Rate4Site scores indicate stronger evolutionary conservation. Population cells report cohort allele frequency and the number of adequately sampled breeds in which the alternative allele was observed. Sites not represented in the source variant dataset, QC-limited estimates, sites at which the alternative allele was not observed, and unavailable or non-assessable molecular evidence are shown separately. Candidate prioritization was based on qualitative convergence across evidence dimensions rather than an arithmetic composite score

The high-priority set comprised NEU3 F159C and E94K, ST8SIA2 R129Q, ST8SIA4 I73L, CMAH N242S and I388M, and ST3GAL4 R203S. NEU3 F159C and E94K, ST8SIA2 R129Q, and ST3GAL4 R203S showed concordance between deleterious SIFT predictions and evolutionary constraint. The remaining high-priority candidates were supported by complementary evidence patterns. ST8SIA4 I73L showed strong evolutionary conservation despite a tolerated SIFT prediction; CMAH N242S combined a deleterious prediction and direct pathway relevance with discordant conservation evidence; and CMAH I388M showed evolutionary conservation and RefSeq-supported protein context despite unavailable native SIFT evidence.

The secondary set comprised CMAH P424L and F512Y, ST8SIA2 S63L, ST8SIA4 R203Q, NEU1 L275P, CMAS E128*, ST3GAL4 *396L, and NANS P302A. These candidates retained biologically relevant support but showed weaker or more discordant convergence across the available evidence dimensions than the high-priority set. Population and breed representation contributed supporting evidence but did not act as an inclusion or exclusion criterion.

### 3.7 Genome-Wide Population-Genomic Analyses

#### 3.7.1 Genome-Wide Population Structure

Genome-wide PCA based on the 950,219 LD-pruned SNPs revealed pronounced genetic structure between South African and European cattle (Figure 7). The first principal component (PC1), explaining approximately 25% of the total genetic variance, separated South African from European breeds, with South African breeds occupying a broader range along PC1. The second principal component (PC2; approximately 11% of variance) further differentiated the European populations, with Finnish, Dutch, and Portuguese breeds showing distinct patterns of distribution. Finnish and Dutch animals formed adjacent but separate clusters, whereas Portuguese animals formed more than one cluster, indicating additional structure within the European cohort. A small number of animals sampled in Portugal clustered with the Dutch breeds (Figure 7B), consistent with the Holstein-Friesian animals sampled there (Table 2).

**Figure 7.**
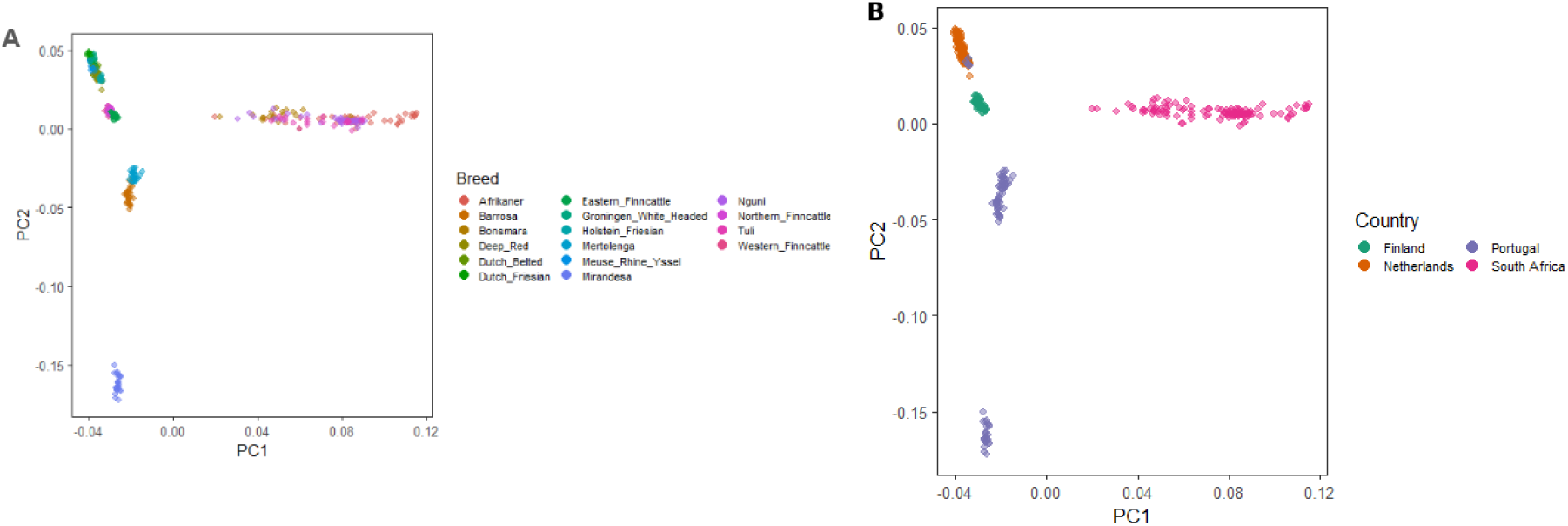
Genome-wide population structure of South African and European cattle. Principal component analysis of the 346 animals based on the 950,219 LD-pruned SNPs, showing PC1 and PC2. (A) Animals colored by breed. (B) Animals colored by sampling country; Holstein-Friesian animals appear under the country in which they were sampled.

#### 3.7.2 Genetic Diversity

Mean nucleotide diversity per ascertained SNP (π) in the 13 sialic-acid pathway genes was highest in the four South African breeds, each of which exceeded all European breeds with at least five animals (Figure 8). Finnish, Dutch, and Portuguese breeds showed lower values, and diversity varied among breeds within each country. Estimates for Mertolenga (n = 2) and Mirandesa (n = 4) rest on very few animals and are interpreted cautiously. Because these estimates are restricted to ascertained pathway SNPs, they describe relative diversity among breeds within the pathway and not genome-wide diversity.

**Figure 8.**
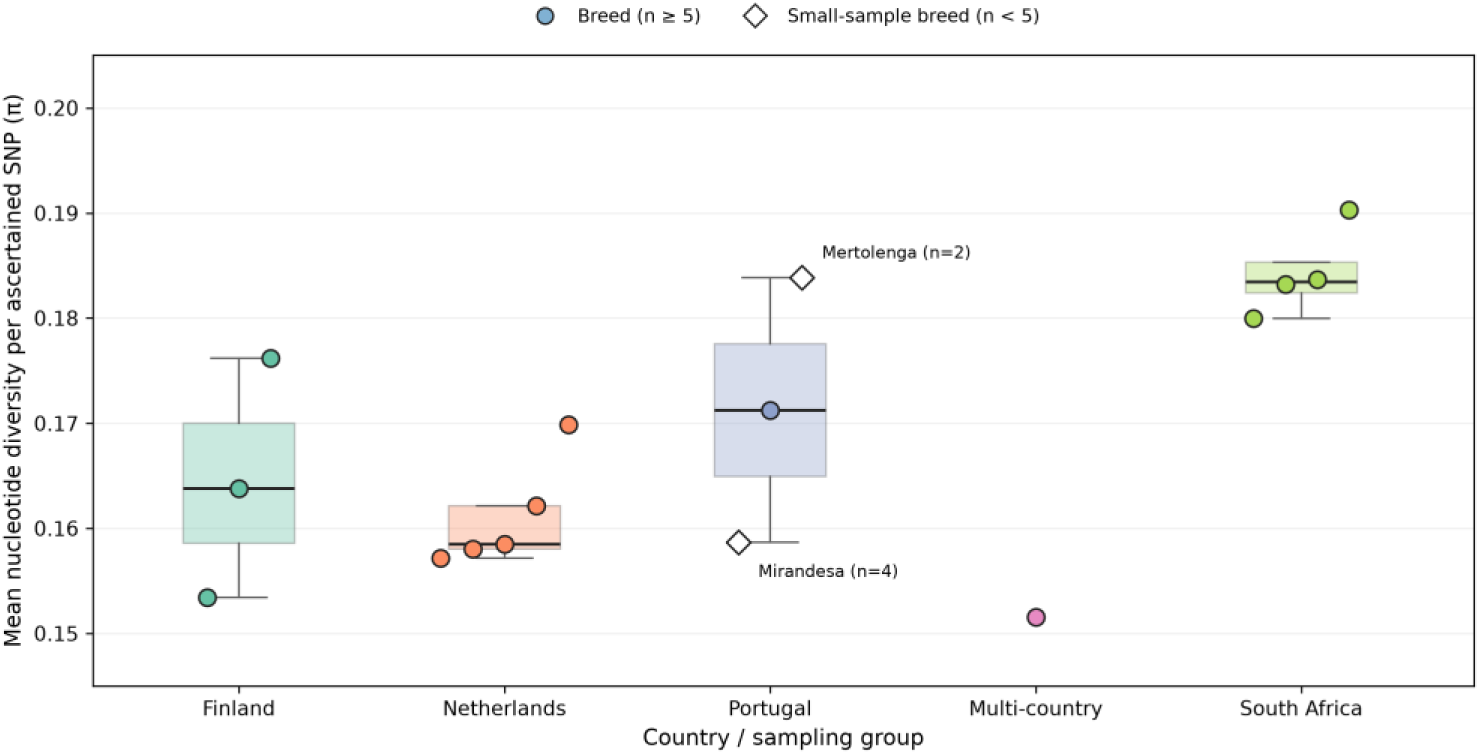
Genetic diversity in the sialic-acid pathway genes of South African and European cattle. Mean nucleotide diversity per ascertained SNP (π) for each breed, calculated from the 4,505 gene-level SNP records and grouped by country of origin. Holstein-Friesian (n = 15), whose animals were sampled in Finland, the Netherlands, and Portugal, is shown as a separate multi-country category. Boxes summarize breeds within each country; open diamonds mark breeds with fewer than five animals (Mertolenga, n = 2; Mirandesa, n = 4). π is calculated per ascertained pathway SNP and is not a genome-wide estimate.

Breed-level distributions of minor allele frequency and of observed and expected heterozygosity for the pathway SNP records are shown in Supplementary Figures S1 and S2. South African breeds had the smallest proportions of monomorphic records (Supplementary Figure S1).

#### 3.7.3 Population Differentiation Across Geographic Scales

Genome-wide scans identified SNPs above the empirical 99.9th-percentile threshold (950 at each level) on all 29 autosomes at every level of comparison, with thresholds rising from the breed-level to the regional scan (0.452, 0.475, and 0.580) (Figure 9A–C).

**Figure 9.**
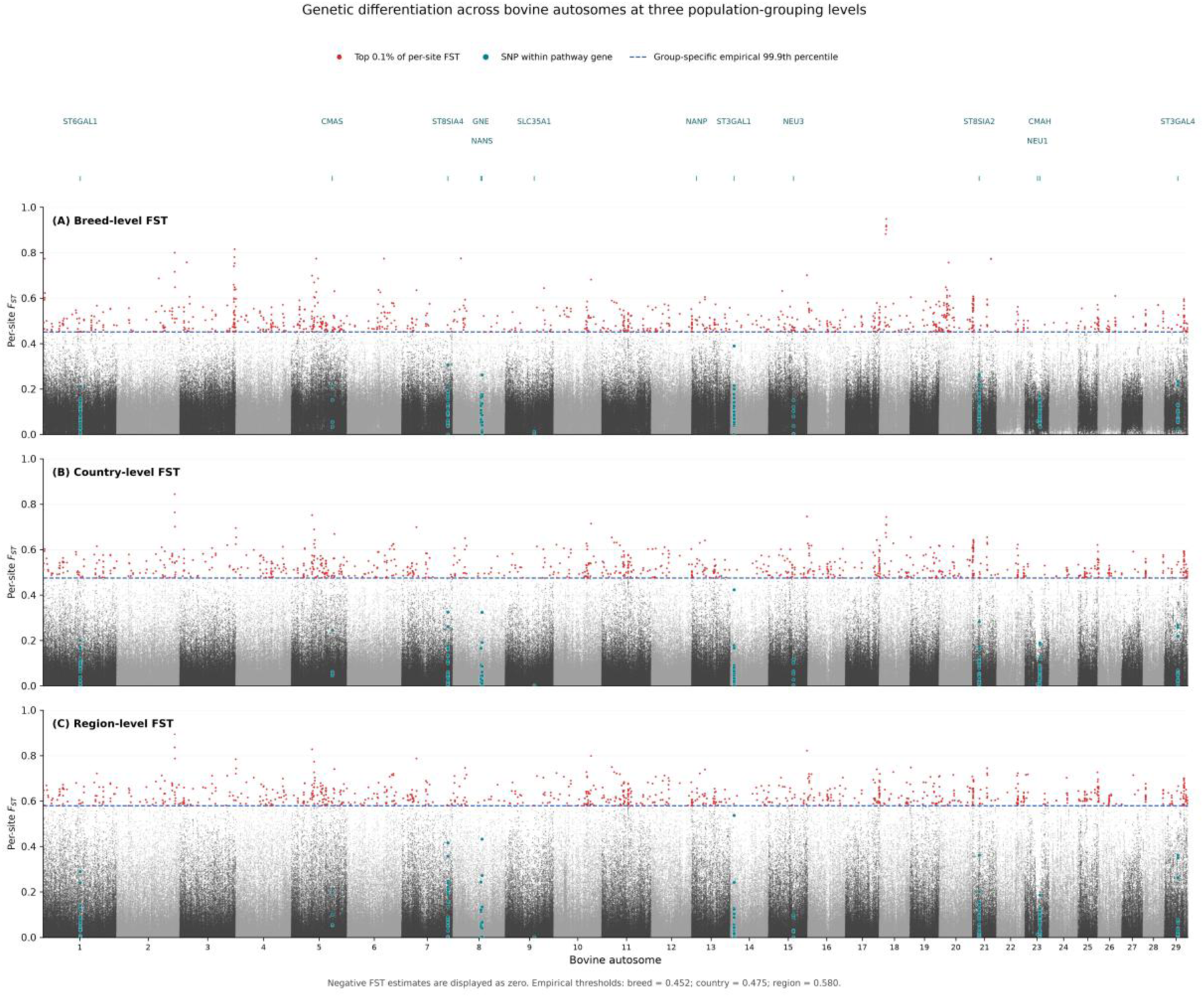
Genetic differentiation across the bovine autosomes at three population-grouping levels. Per-SNP Weir and Cockerham F_ST_ for the 950,219 LD-pruned SNPs at (A) breed, (B) country, and (C) regional level. SNPs within the 13 sialic-acid pathway genes, including the 2-kb upstream extension, are shown in teal, and gene positions are marked above panel A; NANP contained no SNP in the LD-pruned set. Red points mark the top 0.1% of per-SNP F_ST_, and blue dashed lines mark the level-specific empirical 99.9th percentile (breed 0.452, country 0.475, region 0.580). Negative estimates are displayed as zero.

Within the 13 sialic-acid pathway genes, SNPs with F_ST_ ≥ 0.25 were identified at breed level in NANS, ST3GAL1, ST8SIA2, and ST8SIA4. At the country level, ST3GAL4 was additionally identified, and in the South Africa-Europe comparison ST6GAL1 was also highlighted. NANS, ST3GAL1, ST3GAL4, ST8SIA2, and ST8SIA4 recurred across more than one level of comparison. These SNPs are described as highly differentiated and are not interpreted as selection signatures.

Of the 950,219 LD-pruned SNPs, 220 were located within the 13 gene regions, and none within NANP. These pathway SNPs fell below the genome-wide outlier thresholds in all three scans; the highest values (0.390, 0.423, and 0.537 at breed, country, and regional level) all occurred at the same ST3GAL1 position (chr14:7,932,660). Their differentiation therefore lies within the range of variation observed across the genome. The 220 pathway SNPs are distinct from the 62 protein-altering candidates, which were analyzed separately (Sections 2.7.4 and 3.8).

#### 3.7.4 Allele-Frequency Spectrum Across the Sialic-Acid Pathway

Tajima’s D calculated across the pathway SNPs differed between regions (Figure 10). Each value represents one gene in one breed (13 genes in 10 European and 4 South African breeds; 130 and 52 estimates, respectively). Values for European breeds were shifted towards positive values (mean 0.990, with the distribution peaking near 1). South African breeds were centered closer to zero (mean 0.028) and included a larger proportion of negative values. Mertolenga and Mirandesa were excluded from this comparison because their sample sizes were very small (Section 2.7.5). Positive values indicate an excess of intermediate-frequency variants relative to neutral expectation, and negative values an excess of rare variants. Because the gene-level datasets contained segregating SNPs only, and because both demographic history and sampling can shape these values, the results are reported as regional differences in the allele-frequency spectrum and are not interpreted as evidence of selection or of a specific demographic history.

**Figure 10.**
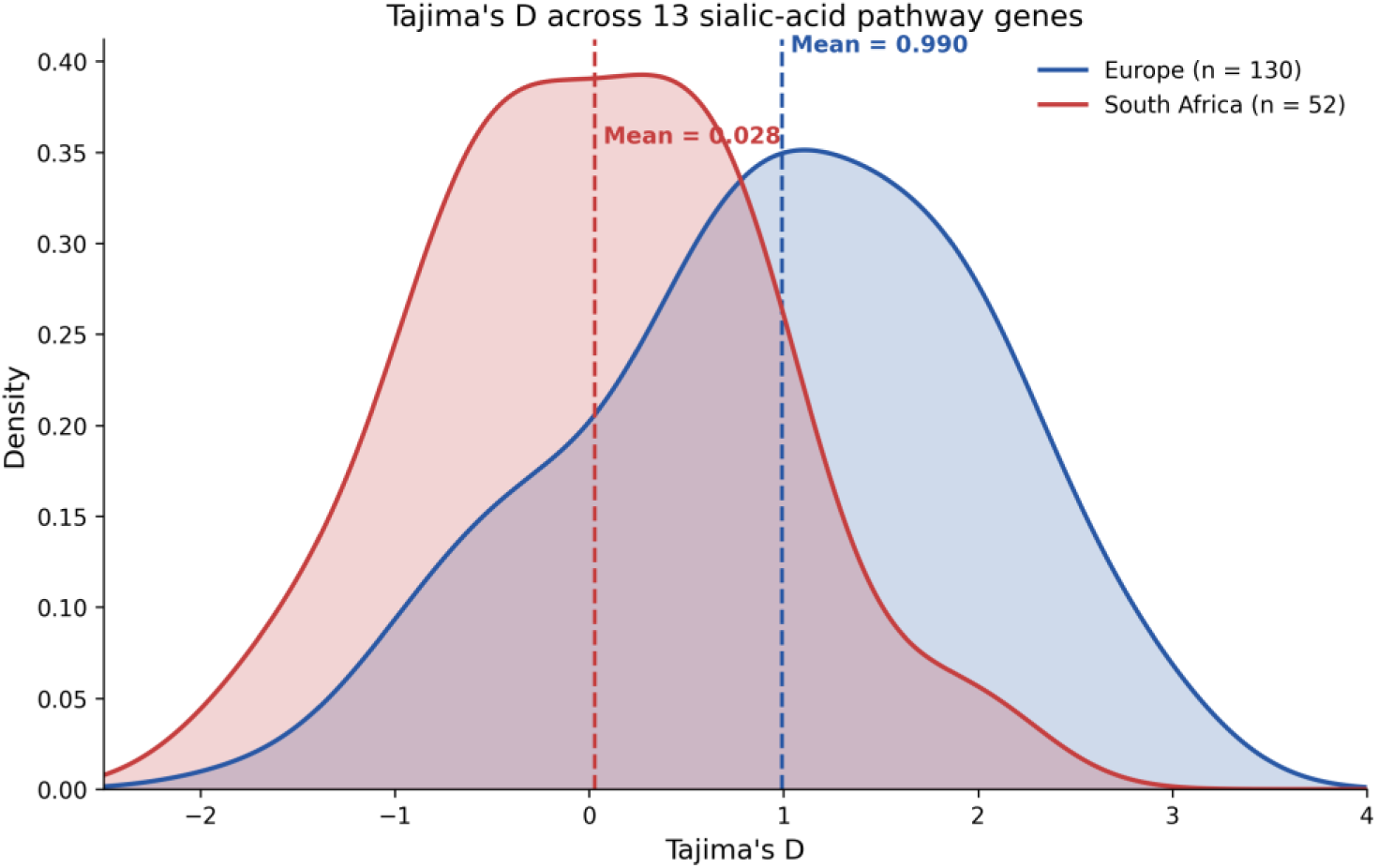
Tajima’s D across the 13 sialic-acid pathway genes in European and South African cattle. Density distributions of breed-by-gene Tajima’s D estimates by region (Europe, 130 estimates from 10 breeds; South Africa, 52 estimates from 4 breeds). Each estimate is the value for one gene in one breed, not for an individual animal. Dashed lines mark the regional means (Europe 0.990; South Africa 0.028). Mertolenga (n = 2) and Mirandesa (n = 4) are excluded because of their small sample sizes.

### 3.8 Integration of Population-Genomic and Functional Evidence

The sialic-acid genes containing SNPs with F_ST_ ≥ 0.25 (Section 3.7.3) were compared with the genes carrying the 15 prioritized candidates (Section 3.6). Three genes occurred in both sets: ST8SIA2 (R129Q, high priority; S63L, secondary), ST8SIA4 (I73L, high priority; R203Q, secondary), and ST3GAL4 (R203S, high priority; *396L, secondary). NANS, which also contained differentiated SNPs, carried one secondary candidate (P302A). ST3GAL1 and ST6GAL1 contained differentiated SNPs but no prioritized candidate. In contrast, NEU3 and CMAH, which together carry four of the seven high-priority variants, were not among the genes with SNPs at F_ST_ ≥ 0.25 in the LD-pruned genome-wide scan. This overlap is at gene level only.

At the variant level, F_ST_ could be estimated for only part of the 62 protein-altering candidates. In the regional comparison, 30 candidates had finite estimates; 21 were absent from the European records, 10 from the South African records, and for one candidate F_ST_ could not be estimated. The highest regional value was observed for the NEU3 variant at chr15:54265634 (F_ST_ = 0.369), followed by NANS chr8:63193083 (0.127) and the ST3GAL4 variants at chr29:29730119, chr29:29730201, and chr29:29730070 (0.084, 0.071, and 0.067). These are the five candidates with the largest regional allele-frequency differences (Section 3.2). Although no NEU3 SNP exceeded F_ST_ ≥ 0.25 in the LD-pruned genome-wide analysis, the candidate-level regional comparison yielded an F_ST_ of 0.369 for chr15:54265634, reflecting the different variant sets and analytical comparisons.

Of the 372 candidate-by-country-pair records (six country pairs), 159 yielded finite estimates. Notable values involved NEU3 in the comparisons of South Africa with the Netherlands (0.442) and Portugal (0.424), and CMAH in the comparisons of Portugal with the Netherlands (0.359) and South Africa (0.319). Because countries differ in breed composition, country-level values may partly reflect the breeds sampled in each country. At breed level, 2,028 of 7,440 candidate-by-breed-pair records (16 breeds, 120 pairs) yielded finite estimates (Supplementary Table S4). Notable values included SLC35A1 in Dutch Belted compared with Mertolenga (0.864) and Mirandesa (0.801), ST3GAL4 in Mertolenga compared with Western Finncattle (0.750), CMAH in Barrosa compared with Dutch Friesian (0.742), and NEU3 in Afrikaner compared with Deep Red (0.683). Three of these five values involve Mertolenga (n = 2) or Mirandesa (n = 4) and are therefore particularly uncertain.

Candidate-level F_ST_ estimates are reported descriptively. High values identify candidates whose allele frequencies differ between the groups compared, but they do not demonstrate selection, functional impact, or a causal association with any phenotype.

Taken together, the genome-wide analyses show that pathway diversity and differentiation follow the population structure of the cohort (Sections 3.7.1 to 3.7.4), and that differentiated pathway SNPs fall within the range of genome-wide variation (Section 3.7.3). Population-specific variation within the sialic-acid pathway therefore occurs against a broader background of genomic differentiation, and allele-frequency differences among the prioritized variants (Section 3.2) were not considered evidence of pathway-specific selection in isolation. The functional prioritization of Section 3.6 identifies individual variants within that background and does not rely on differentiation statistics.

## 4. Discussion

This study provides a variant-level population-genomic analysis of 13 genes spanning the bovine sialic acid network in European and South African cattle. Earlier work on the genetics of bovine sialic acid biology addressed single genes, such as CMAH (Ogun et al., 2023a), or milk oligosaccharide traits through genome-wide association analysis (Poulsen et al., 2019). Across the 13 genes, we identified 62 protein-altering candidates and prioritized 15 using functional, evolutionary and protein-context evidence, with population representation as supporting context. Most target-gene variation was noncoding: 8,675 of the 9,724 variants were intronic, so the protein-altering subset is a small part of the variation in this network. Within that subset, population differentiation and functional evidence nominated different variants. We first describe this separation, then consider how the candidates were prioritized, their biological context, the population background in which they occur and the implications of their rarity for follow-up studies.

The candidates that differed most between populations were not those with the strongest functional evidence. NEU3 Y136H (chr15:54265634) showed the largest regional contrast, with an alternative-allele frequency of 0.83 in South African and 0.34 in European cattle (ΔAF = 0.48), and the highest regional candidate-level F_ST_ (0.369). NANS H16Q and the ST3GAL4 substitutions L59F, R87C and P43L completed the five most differentiated candidates. None of the five was prioritized, although ST3GAL4 P43L carried a low-confidence deleterious SIFT prediction. Conversely, many prioritized candidates were rare. ST8SIA2 R129Q, NEU3 E94K, CMAH I388M, CMAH P424L and ST3GAL4 *396L were each represented by a single alternative allele in the South African cohort, and CMAH N242S by one South African and six European alleles. ST3GAL4 R203S was prioritized on molecular evidence, but its alternative allele was not observed in the analyzed South African genotypes and the site was absent from the European data, so its segregation in these cohorts remains unconfirmed. ST8SIA4 I73L was the clearest exception, occurring in seven European breeds at an overall European allele frequency of 0.096.

This separation follows from what the two kinds of evidence measure. F_ST_ summarizes allele-frequency divergence between groups, whereas SIFT and Rate4Site estimate whether a substitution affects a conserved residue. F_ST_ is also tightly constrained when the minor allele is rare, so sites with rare minor alleles yield small values (Jakobsson et al., 2013). Rare substitutions are therefore unlikely to appear among the most differentiated variants, whatever their functional evidence. The gene-level comparison is consistent with this. ST8SIA2, ST8SIA4 and ST3GAL4 contained both highly differentiated pathway SNPs and prioritized candidates, but the overlap is at gene level rather than at the same variant. NEU3 and CMAH, which together carry four of the seven high-priority candidates, contained no pathway SNP at F_ST_ ≥ 0.25 in the genome-wide scan. Candidate-level and genome-wide F_ST_ were estimated on different variant sets with different filters, so genome-wide thresholds do not transfer to candidates. Weir and Cockerham estimates can also vary with the ratio of sample sizes between groups (Bhatia et al., 2013), and ΔAF and F_ST_ describe the same allele-frequency pattern rather than independent evidence. NEU3 Y136H is therefore best regarded as a population-differentiated candidate. Differentiation and functional prioritization are complementary: one locates allele-frequency contrasts and the other locates plausible protein changes, and in this dataset no candidate clearly satisfied both.

Within the prioritized set, both convergence and discordance were informative. For ST8SIA2 R129Q, NEU3 E94K and F159C, and ST3GAL4 R203S, deleterious SIFT predictions coincided with evolutionary constraint and residue-level context. Because SIFT itself uses sequence homology (Ng and Henikoff, 2003), this agreement represents complementary, partly overlapping evidence rather than independent confirmation. ST8SIA4 I73L had the strongest conservation signal among the candidates despite a tolerated SIFT prediction, whereas CMAH N242S combined a deleterious prediction with discordant conservation. CMAH I388M lacked a native SIFT annotation and was retained on the basis of conservation, exact RefSeq support and pathway relevance. For the non-missense candidates, the stop-gained annotation for CMAS E128* and the stop-lost annotation for ST3GAL4 *396L describe the change in coding sequence but not its consequence for the protein. Retaining these discordant and incomplete cases, rather than averaging them into a score, identifies the variants for which experimental data would be most decisive.

The prioritized candidates lie in genes that control Neu5Gc formation, polysialylation, sialic acid turnover and terminal sialylation. CMAH converts CMP-Neu5Ac to CMP-Neu5Gc (Chou et al., 1998). It carried two high-priority candidates, N242S and I388M, and two secondary candidates, P424L and F512Y, while CMAH K107E varied markedly in frequency among European breeds. Neu5Gc content varies among cattle breeds in both meat and milk. In beef, significant differences were reported among seven breeds, particularly for Neu5Gc (Contò et al., 2025). In milk, Neu5Gc was higher in Holstein, Simmental and their crosses than in Podolica. Podolica was kept in an extensive grazing system and the other breeds were housed intensively, so breed and management cannot be separated (Crisà et al., 2022). Neu5Gc is also present on milk oligosaccharides in early lactation, with wide differences between cows (Isernhagen et al., 2024). These observations document breed-level and individual variation in bovine Neu5Gc, and the CMAH candidates make its genetic contribution testable. The question extends beyond animal biology. Dietary Neu5Gc is metabolically incorporated into human tissues, and in Neu5Gc-deficient mice, dietary Neu5Gc combined with anti-Neu5Gc antibodies promoted inflammation and tumor progression (Samraj et al., 2015).

The remaining candidates occupy equally relevant steps. ST8SIA2 R129Q and ST8SIA4 I73L lie in the two polysialyltransferases. Both genes are expressed in bovine udder tissue and in cultivated mammary epithelial cells, which are a source of milk polySia, and milk polySia declines during the first three days of lactation in line with Neu5Ac (Hinterseher et al., 2023). NEU3, cloned and characterized from bovine brain, is a plasma membrane-associated sialidase that hydrolyzes gangliosides almost specifically (Miyagi et al., 1999). Human ST3GAL4 transfers sialic acid in α2,3-linkage to glycoprotein and glycolipid acceptors (Kitagawa and Paulson, 1994), and in mice it is required for L-selectin-dependent leukocyte rolling during inflammation (Sperandio et al., 2006). In bovine milk, 3′-sialyllactose falls by more than half within one day of calving (Isernhagen et al., 2024). These functions place the candidates at steps that shape the sialoglycome. Whether the bovine substitutions alter enzyme activity or milk glycan composition requires direct testing.

The population background helps interpret where these candidates occur. Diversity per ascertained pathway SNP was higher in all four South African breeds than in the European breeds with at least five animals, and South African breeds had the smallest proportions of monomorphic records. This agrees with their origins. Sanga breeds are composites of African taurine and Bos indicus cattle, and Bonsmara was developed by crossbreeding Afrikaner with Hereford and Shorthorn (Makina et al., 2016). It also agrees with the high genetic diversity reported for African zebu and Sanga cattle from other regions of the continent (Kim et al., 2017). Our PCA describes population structure rather than ancestry proportions, so admixture is a plausible contributor rather than a demonstrated cause.

Tajima’s D showed a regional pattern, shifted toward positive values in European breeds and centered near zero in South African breeds. Positive values indicate an excess of intermediate-frequency variants and can result from population bottlenecks as well as balancing selection (Akey et al., 2004). Stepwise reductions in effective population size have been inferred for Holstein (MacLeod et al., 2013), one of the ten European breeds. Data assembly offers a second explanation. The European and South African datasets were generated separately, and the diversity and frequency-spectrum analyses used only SNP records present in both, so region is partly confounded with cohort. Under-representation of rare variants also biases Tajima’s D upward (Clark et al., 2005). For the same reason, our diversity values compare breeds within the ascertained SNP set and are not per-base estimates, which require invariant and missing sites to be accounted for (Korunes and Samuk, 2021). We therefore report the regional contrast descriptively. The differentiation result was clear: none of the 220 LD-pruned pathway SNPs exceeded the genome-wide thresholds at breed, country or regional level. The network, as represented by these markers, followed the genomic background of the two cohorts.

The rarity of the prioritized candidates has direct consequences for study design. In Danish Holstein and Jersey cattle, acidic milk oligosaccharides had low to insignificant heritabilities, and no significant SNPs were found for 3′-sialyllactose, 6′-sialyllactose or 2 Hex 2 NeuAc (Poulsen et al., 2019). That study addressed variation captured in two breeds, and its results neither include nor exclude rare variants of the kind prioritized here. With one or a few carriers, such variants cannot be evaluated by association in cohorts of this size. Their study requires carriers to be identified by targeted genotyping and then compared for enzyme function or milk composition. Single observations also need independent confirmation, and absence from a variant-only dataset does not establish absence from a breed.

These results should be interpreted in light of the study design. The analysis combined two independently generated whole-genome datasets covering 16 breeds and three grouping levels, which allowed the pathway to be examined against a genome-wide background, although breed representation was uneven, with few animals for Mertolenga and Mirandesa. The functional evidence was computational, so the prioritized set defines testable hypotheses for experimental work. Variants were called against the taurine ARS-UCD1.2 assembly, and because a breed-diverse graph genome improved read mapping most for the indicine Sahiwal breed (Talenti et al., 2022), some divergent sequence may be under-represented in indicine-influenced breeds. Regulatory effects of noncoding variation and matched milk phenotypes were beyond the scope of this study and are natural extensions, supported by emerging regulatory atlases for cattle (Liu et al., 2022).

These results define complementary next steps. Targeted genotyping in larger and more diverse cohorts would confirm single observations and establish carrier frequencies. Combined with haplotype analysis, it would show whether prioritized and differentiated variants within the same genes share haplotype backgrounds. In parallel, wild-type and variant enzymes can be compared in assays appropriate to each protein to determine whether the substitutions alter activity, substrate preference or localization. Genotype-informed measurement of Neu5Ac, Neu5Gc, polySia and sialylated oligosaccharides in milk would then test for associations with the milk sialoglycome. These compounds change within days of calving, and individual differences may reflect metabolic stress during the transition period (Isernhagen et al., 2024). Such studies therefore require precisely timed early-lactation sampling and records of physiological state. Harmonized data generation that retains callable-site information would also allow per-base diversity estimates and help separate population differences from cohort effects.

## 5. Conclusions

This study identified 62 protein-altering candidates across 13 genes of the bovine sialic acid network in 346 European and South African cattle and prioritized 15 using functional, evolutionary and protein-context evidence. Population differentiation and functional prioritization nominated different variants. NEU3 Y136H was the most differentiated candidate but was not prioritized, while many prioritized candidates were rare.

The 220 pathway markers fell within the genome-wide range of differentiation, and diversity within the ascertained SNP set was higher in the South African breeds, consistent with their composite origins. These analyses do not test selection or establish phenotypic effects.

The candidate set provides a basis for targeted genotyping, enzyme-level testing and genotype-informed early-lactation milk phenotyping. Differentiation alone did not identify the candidates with the strongest functional evidence, which supports combining population and molecular evidence when selecting variants for experimental follow-up.

## Data Availability

Whole-genome sequencing variant data for European and South African cattle were obtained from Ginja et al. (2025) and Dlamini et al. (2026), respectively. The European cattle variant dataset is available in the European Variation Archive (EVA) under accession PRJEB98152, and the South African cattle variant datasets are available under accessions PRJEB110435 and PRJEB94226. Raw sequencing reads are available in the European Nucleotide Archive (ENA) under accessions PRJEB90816 and PRJEB90914 for European and South African cattle, respectively. All analysis code, workflow configurations, environment specifications, and variant-annotation scripts will be deposited in the project’s GitHub repository (https://github.com/Mayorcool-lab/bovine-sialic-acid-popgen) and archived on Zenodo with a persistent DOI upon manuscript submission.

## Authors’ contributions

O.J.O. conceived and designed the study, obtained and curated the public datasets, performed variant annotation, functional prioritization and pathway- and candidate-level population-genomic analyses, validated and revised the initial genome-wide analyses, prepared the figures and tables, interpreted the results and drafted the manuscript. L.V.A. performed the initial genome-wide principal component, differentiation, diversity and Tajima’s D analyses. D.B. advised on genomic resources, dataset selection, variant annotation strategy and data quality control. S.P.G. contributed to the glycobiological interpretation and biological framing of the sialic acid network and milk sialoglycome and critically reviewed the manuscript. All authors read and approved the final manuscript.

## Conflicts of Interest

The authors declare no conflicts of interest.

## Acknowledgements

The authors thank the FAANG consortium for making bovine functional genomic annotations available. The precise wording of the acknowledgement for genomic data access and computational resources will be completed once the datasets and computing infrastructure have been confirmed. [O.J.O.: add any specific acknowledgements for computational resources used.]

## Supplementary Materials

**Supplementary Table S1**. Breed-level allele frequencies of the 62 protein-altering candidate variants across 16 South African and European cattle breeds.

Allele frequencies are reported for 62 protein-altering candidate variants across four South African and 12 European cattle breeds. Status indicates whether a variant was present, absent after site-level quality control, or not represented in the source variant dataset. Estimates based on fewer than five callable animals are flagged as very small samples.

**Supplementary Table S2.** Protein and functional context of the 62 protein-altering candidate variants. For each candidate, the table reports the mapping between the variant reference protein and the annotated protein, overlap with UniProt and InterPro features, and the resulting context level (residue-level or protein-level only). ST3GAL4 candidates were reconciled across multiple reference proteins, and RefSeq-resolved CMAH variants were annotated by sequence-based analysis.

**Supplementary Table S3.** Integrated prioritization of 15 protein-altering candidates across the bovine sialic-acid network. Seven candidates were classified as high priority and eight as secondary priority based on qualitative convergence across functional prediction, evolutionary conservation, protein context, pathway relevance, and population evidence. Missing or non-assessable evidence was not treated as negative, and no arithmetic composite score was applied.

| Candidate | Consequence | SIFT | Rate4Site | Population evidence | Priority | Key interpretation |
|---|---|---|---|---|---|---|
| NEU3 F159C | Missense | Deleterious (0.01) | -0.5026 | SA: AF 0.0084 (1 breed); EU: AF 0.0090 (3 breeds) | High priority | Concordant deleterious SIFT, evolutionary conservation and residue-level context; ALT observed in both populations, although current allele frequencies are low. |
| ST8SIA4 I73L | Missense | Tolerated (0.54) | -0.9582 | SA: AF 0.0087 (2 breeds); EU: AF 0.0962 (7 breeds) | High priority | Strongest conservation signal among the prioritised candidates and broad population segregation despite a tolerated SIFT prediction. |
| ST8SIA2 R129Q | Missense | Deleterious (0.01) | -0.9269 | SA: AF 0.0043 (1 breed); EU: not in source | High priority | Deleterious SIFT, strong evolutionary conservation and residue-level context; current population evidence is limited to South Africa. |
| NEU3 E94K | Missense | Deleterious (0) | -0.6812 | SA: AF 0.0043 (1 breed); EU: not in source | High priority | Concordant deleterious SIFT, evolutionary conservation and residue-level context; currently observed in one South African breed. |
| CMAH N242S | Missense | Deleterious (0.03) | 0.3811 | SA: AF 0.0042 (1 breed); EU: AF 0.0136 (3 breeds) | High priority | Deleterious SIFT, residue-level context and direct relevance to Neu5Gc biosynthesis, but conservation evidence is discordant. |
| ST3GAL4 R203S | Missense | Deleterious (0.01) | -0.2956 | SA: ALT not observed; EU: not in source | High priority | Deleterious SIFT and evolutionary conservation with robust multi-reference residue reconciliation; current ALT representation is limited. |
| CMAH I388M | Missense | Unavailable | -0.6392 | SA: AF 0.0043 (1 breed); EU: not in source | High priority | Evolutionary conservation, exact RefSeq-X2 support and direct CMAH pathway relevance despite unavailable native SIFT evidence. |
| CMAH P424L | Missense | Unavailable | -0.5806 | SA: AF 0.0046 (1 breed); EU: ALT not observed | Secondary priority | Conservation, exact RefSeq-X2 support and direct CMAH pathway relevance, but with weaker conservation than I388M and unavailable native SIFT evidence. |
| CMAH F512Y | Missense | Unavailable | -0.0075 | SA: QC-limited AF 0.0510; EU: AF 0.0498 (6 breeds) | Secondary priority | Direct CMAH pathway relevance, exact RefSeq-X2 support and broad European segregation, but little conservation support and unavailable native SIFT evidence. |
| ST8SIA2 S63L | Missense | Tolerated (0.07) | -0.5438 | SA: not in source; EU: AF 0.0091 (1 breed) | Secondary priority | Evolutionary conservation and residue-level context despite a tolerated SIFT prediction; current population support is limited. |
| ST8SIA4 R203Q | Missense | Tolerated (0.6) | -0.8719 | SA: not in source; EU: ALT not observed | Secondary priority | Strong evolutionary conservation and residue-level context despite a tolerated SIFT prediction; current ALT representation is limited. |
| NEU1 L275P | Missense + splice region | Deleterious (0) | Not assessable | SA: AF 0.0084 (2 breeds); EU: AF 0.0294 (4 breeds) | Secondary priority | Deleterious SIFT and combined missense/splice-region consequence with ALT observed in both populations, but conservation was not assessable and context is protein-level only. |
| CMAS E128* | Stop gained | Not applicable | -0.8396 | SA: QC-limited AF 0.0093; EU: ALT not observed | Secondary priority | Stop-gained consequence with strong evolutionary conservation; South African evidence is QC-limited and ALT was not observed in the European cohort. |
| ST3GAL4 *396L | Stop lost | Not applicable | Special case | SA: AF 0.0042 (1 breed); EU: not in source | Secondary priority | Distinct stop-lost consequence with ALT observed in one South African breed; conservation was treated as a special case and context is protein-level only. |
| NANS P302A | Missense | Tolerated (0.16) | -0.5498 | SA: QC-limited AF 0.0429; EU: AF 0.0091 (2 breeds) | Secondary priority | Evolutionary conservation and residue-level context despite a tolerated SIFT prediction; South African evidence is QC-limited. |

*Note: QC-limited allele frequencies were calculated from callable genotypes at sites that did not meet the standard site-level call-rate threshold and should therefore be interpreted cautiously. “Not in source” indicates that the site was not represented in the source variant dataset and no allele frequency was assigned.*

**Supplementary Table S4.** Pairwise breed-level F_ST_ for the 62 protein-altering candidate variants. Weir and Cockerham F_ST_ for 7,440 candidate-by-breed-pair records (16 breeds, 120 pairs), of which 2,028 yielded finite estimates. Candidates absent from the variant-only VCF records of a breed were not imputed as reference genotypes, and the corresponding comparisons are non-estimable. Raw negative estimates are retained. Comparisons involving Mertolenga (n = 2) or Mirandesa (n = 4) should be interpreted with particular caution.

**Supplementary Figure S1.** Breed-specific minor allele frequency distribution across the 4,505 ascertained pathway SNP records. For each breed, records are classified as monomorphic (MAF = 0), low-frequency (0 < MAF < 0.05), common (MAF ≥ 0.05), or uncallable (MAF unavailable). Mertolenga (n = 2) and Mirandesa (n = 4) are marked as small samples, and their distributions are strongly affected by sample size.

**Supplementary Figure S2.** Observed (H_O_) and expected (H_E_) heterozygosity by breed across the 4,505 ascertained pathway SNP records. H_O_ is the proportion of heterozygous genotypes among called genotypes, and H_E_ is the mean of 2p(1 − p) across records (Section 2.7.3). Estimates describe the ascertained pathway records only and are not genome-wide values. Mertolenga (n = 2) and Mirandesa (n = 4) are marked as small samples.

## Notes

### Competing Interest Statement

The authors have declared no competing interest.

### Summary of Updates

Dear Editorial Team, Thank you for making this article available on your reputable platform. However, there are two errors on the name of the authors. These two authors do not have "PA" in their names. Firstly, Peter A. Idowu and Doreen Becker. Please write the name as seen on the manuscript as attached in this email We hope you can assist with this. Thank you so much for your assistance Regards

## References

Akey, J.M., Eberle, M.A., Rieder, M.J., Carlson, C.S., Shriver, M.D., Nickerson, D.A., Kruglyak, L. (2004). Population history and natural selection shape patterns of genetic variation in 132 genes. PLoS Biology, 2(10), e286. DOI: 10.1371/journal.pbio.0020286.

Ashkenazy, H., Abadi, S., Martz, E., Chay, O., Mayrose, I., Pupko, T., Ben-Tal, N. (2016) ConSurf 2016: an improved methodology to estimate and visualize evolutionary conservation in macromolecules. Nucleic Acids Research, 44, W344–W350. DOI: 10.1093/nar/gkw408.

Bhatia, G., Patterson, N., Sankararaman, S., Price, A.L. (2013). Estimating and interpreting FST: the impact of rare variants. Genome Research, 23(9), 1514–1521. DOI: 10.1101/gr.154831.113.

Chang, C.C., Chow, C.C., Tellier, L.C.A.M., Vattikuti, S., Purcell, S.M., Lee, J.J. (2015) Second-generation PLINK: rising to the challenge of larger and richer datasets. GigaScience, 4, 7. DOI: 10.1186/s13742-015-0047-8.

Chou, H.-H., Takematsu, H., Diaz, S., Iber, J., Nickerson, E., Wright, K.L., Muchmore, E.A., Nelson, D.L., Warren, S.T., Varki, A. (1998). A mutation in human CMP-sialic acid hydroxylase occurred after the Homo-Pan divergence. Proceedings of the National Academy of Sciences of the United States of America, 95(20), 11751–11756. DOI: 10.1073/pnas.95.20.11751.

Clark, A.G., Hubisz, M.J., Bustamante, C.D., Williamson, S.H., Nielsen, R. (2005). Ascertainment bias in studies of human genome-wide polymorphism. Genome Research, 15(11), 1496–1502. DOI: 10.1101/gr.4107905.

Contò, M., Miarelli, M., Di Giovanni, S., Failla, S. (2025). Variability of sialic acids in beef breeds and nutritional implications in red meat. Molecules, 30(3), 710. DOI: 10.3390/molecules30030710.

Crisà, A., Marchitelli, C., Failla, S., Contò, M. (2022). Determination of N-acetylneuraminic and N-glycolylneuraminic acids in unprocessed milk of four cattle breeds. Journal of Dairy Research, 89, 299–301. DOI: 10.1017/S0022029922000620.

Danecek, P., Auton, A., Abecasis, G., Albers, C.A., Banks, E., DePristo, M.A., Handsaker, R.E., Lunter, G., Marth, G.T., Sherry, S.T., McVean, G., Durbin, R. (2011) The variant call format and VCFtools. Bioinformatics, 27, 2156–2158. DOI: 10.1093/bioinformatics/btr330.

Danecek, P., Bonfield, J.K., Liddle, J., Marshall, J., Ohan, V., Pollard, M.O., Whitwham, A., Keane, T., McCarthy, S.A., Davies, R.M., Li, H. (2021) Twelve years of SAMtools and BCFtools. GigaScience, 10, giab008. DOI: 10.1093/gigascience/giab008.

Dhar, C., Sasmal, A., Varki, A. (2019). From “serum sickness” to “xenosialitis”: Past, present, and future significance of the non-human sialic acid Neu5Gc. Frontiers in Immunology, 10, 807. DOI: 10.3389/fimmu.2019.00807.

Dlamini, N., Gao, J., Ginja, C., Kantanen, J., Ghanem, N., Kugonza, D.R., Makgahlela, M., Elnahas, A., Zwane, A., Zawedde, B.M., Nakkazi, C., Behabura, G., Nxumalo, K., Malima, M., Radwan, M.A., Elsawy, M.H., Agaba, M., Fahim, N.H., Khfagy, R.A., Agamy, R., Okwasiimire, R., Waibi, S., Lashmar, S., Crooijmans, R.P.M.A. (2026). Whole-genome sequences of 240 indigenous African cattle from Egypt, Uganda, and South Africa. Scientific Data, 13, 906. DOI: 10.1038/s41597-026-07458-y.

Eisenberg, I., Avidan, N., Potikha, T., Hochner, H., Chen, M., Olender, T., Barash, M., Shemesh, M., Sadeh, M., Grabov-Nardini, G., Shmilevich, I., Friedmann, A., Karpati, G., Bradley, W.G., Baumbach, L., Lancet, D., Asher, E.B., Beckmann, J.S., Argov, Z., Mitrani-Rosenbaum, S. (2001). The UDP-N-acetylglucosamine 2-epimerase/N-acetylmannosamine kinase gene is mutated in recessive hereditary inclusion body myopathy. Nature Genetics, 29, 83–87. DOI: 10.1038/ng718.

Flynn, R.A., Pedram, K., Malaker, S.A., Batista, P.J., Smith, B.A.H., Johnson, A.G., George, B.M., Majzoub, K., Villalta, P.W., Carette, J.E., Bertozzi, C.R. (2021). Small RNAs are modified with N-glycans and displayed on the surface of living cells. Cell, 184(12), 3109–3124.e22. DOI: 10.1016/j.cell.2021.04.023.

Ginja, C., Gao, J., Kantanen, J., Ghanem, N., Kugonza, D., Makgahlela, M., Pires, A.E., Usié, A., Dibbits, B., de Sousa, C.B., Gaspar, D., Ruvinskiy, D., Bijl, E., Smidt, H., Lindeberg, H., Bovenhuis, H., Laport, K., Pokharel, K., Blaschikoff, L., Weldenegodguad, M., Gonzalez Prendes, R., Okwasiimire, R., Guimarães, S., Lui, Y., Crooijmans, R.P.M.A. (2025). Whole genome sequences of 289 native cattle from Finland, the Netherlands, and Portugal. Scientific Data, 12, 1897. DOI: 10.1038/s41597-025-06188-x.

Herrero, J., Muffato, M., Beal, K., Fitzgerald, S., Gordon, L., Pignatelli, M., Vilella, A.J., Searle, S.M.J., Amode, R., Brent, S., Spooner, W., Kulesha, E., Yates, A., Flicek, P. (2016) Ensembl comparative genomics resources. Database, 2016, bav096. DOI: 10.1093/database/bav096.

Hinterseher, J., Günther, J., Zlatina, K., Isernhagen, L., Viergutz, T., Wirthgen, E., Hoeflich, A., Vernunft, A., Galuska, S.P. (2023). Milk polysialic acid levels rapidly decrease in line with the N-acetylneuraminic acid concentrations during early lactation in dairy cows. Biology, 12(1), 5. DOI: 10.3390/biology12010005.

Isernhagen, L., Galuska, C.E., Vernunft, A., Galuska, S.P. (2024). Structural characterization and abundance of sialylated milk oligosaccharides in Holstein cows during early lactation. Foods, 13(16), 2484. DOI: 10.3390/foods13162484.

Isernhagen, L., Galuska, C.E., Hoeflich, A., Galuska, S.P. (2026a). Analysis of the milk oligosaccharides spectrum and sialylation status of milk from West African Dwarf goat and East Friesian sheep. ACS Omega, 11, 32310–32320. DOI: 10.1021/acsomega.5c13396.

Isernhagen, L., Helffenstein, K., Seidel, A., Kuhla, B., Galuska, S.P. (2026b). Monitoring of the sialylation and polysialylation status of sheep milk during early lactation. JDS Communications, 7(5), 547–552. DOI: 10.3168/jdsc.2026-1050.

Jakobsson, M., Edge, M.D., Rosenberg, N.A. (2013). The relationship between FST and the frequency of the most frequent allele. Genetics, 193(2), 515–528. DOI: 10.1534/genetics.112.144758.

Katoh, K., Standley, D.M. (2013) MAFFT multiple sequence alignment software version 7: improvements in performance and usability. Molecular Biology and Evolution, 30, 772–780. DOI: 10.1093/molbev/mst010.

Kim, J., Hanotte, O., Mwai, O.A., Dessie, T., Bashir, S., Diallo, B., Agaba, M., Kim, K., Kwak, W., Sung, S., Seo, M., Jeong, H., Kwon, T., Taye, M., Song, K.-D., Lim, D., Cho, S., Lee, H.-J., Yoon, D., Oh, S.J., Kemp, S., Lee, H.-K., Kim, H. (2017). The genome landscape of indigenous African cattle. Genome Biology, 18, 34. DOI: 10.1186/s13059-017-1153-y.

Kitagawa, H., Paulson, J.C. (1994). Cloning of a novel α2,3-sialyltransferase that sialylates glycoprotein and glycolipid carbohydrate groups. Journal of Biological Chemistry, 269(2), 1394–1401. PMID: 8288606.

Korunes, K.L., Samuk, K. (2021). pixy: Unbiased estimation of nucleotide diversity and divergence in the presence of missing data. Molecular Ecology Resources, 21(4), 1359–1368. DOI: 10.1111/1755-0998.13326.

Lasekan, J., Choe, Y., Dvoretskiy, S., Devitt, A., Zhang, S., Mackey, A., Wulf, K., Buck, R., Steele, C., Johnson, M., Baggs, G. (2022). Growth and gastrointestinal tolerance in healthy term infants fed milk-based infant formula supplemented with five human milk oligosaccharides (HMOs): A randomized multicenter trial. Nutrients, 14(13), 2625. DOI: 10.3390/nu14132625.

Li, C., Liu, Z., Li, M., Miao, M., Zhang, T. (2025). Review on bioproduction of sialylated human milk oligosaccharides: Synthesis methods, physiologic functions, and applications. Carbohydrate Polymers, 352, 123177. DOI: 10.1016/j.carbpol.2024.123177.

Liu, S., Gao, Y., Canela-Xandri, O., Wang, S., Yu, Y., Cai, W., Li, B., Xiang, R., Chamberlain, A.J., Pairo-Castineira, E., D’Mellow, K., Rawlik, K., Xia, C., Yao, Y., Navarro, P., Rocha, D., Li, X., Yan, Z., Li, C., Rosen, B.D., Van Tassell, C.P., Vanraden, P.M., Zhang, S., Ma, L., Cole, J.B., Liu, G.E., Tenesa, A., Fang, L. (2022). A multi-tissue atlas of regulatory variants in cattle. Nature Genetics, 54, 1438–1447. DOI: 10.1038/s41588-022-01153-5.

MacLeod, I.M., Larkin, D.M., Lewin, H.A., Hayes, B.J., Goddard, M.E. (2013). Inferring demography from runs of homozygosity in whole-genome sequence, with correction for sequence errors. Molecular Biology and Evolution, 30(9), 2209–2223. DOI: 10.1093/molbev/mst125.

Makina, S.O., Whitacre, L.K., Decker, J.E., Taylor, J.F., MacNeil, M.D., Scholtz, M.M., van Marle-Köster, E., Muchadeyi, F.C., Makgahlela, M.L., Maiwashe, A. (2016). Insight into the genetic composition of South African Sanga cattle using SNP data from cattle breeds worldwide. Genetics Selection Evolution, 48, 88. DOI: 10.1186/s12711-016-0266-1.

Martín, M.J., Martín-Sosa, S., García-Pardo, L.A., Hueso, P. (2001). Distribution of bovine milk sialoglycoconjugates during lactation. Journal of Dairy Science, 84(5), 995–1000. DOI: 10.3168/jds.S0022-0302(01)74558-4.

McLaren, W., Gil, L., Hunt, S.E., Riat, H.S., Ritchie, G.R.S., Thormann, A., Flicek, P., Cunningham, F. (2016) The Ensembl Variant Effect Predictor. Genome Biology, 17, 122. DOI: 10.1186/s13059-016-0974-4.

Miyagi, T., Wada, T., Iwamatsu, A., Hata, K., Yoshikawa, Y., Tokuyama, S., Sawada, M. (1999). Molecular cloning and characterization of a plasma membrane-associated sialidase specific for gangliosides. Journal of Biological Chemistry, 274(8), 5004–5011. DOI: 10.1074/jbc.274.8.5004.

Ng, P.C., Henikoff, S. (2003) SIFT: predicting amino acid changes that affect protein function. Nucleic Acids Research, 31, 3812–3814. DOI: 10.1093/nar/gkg509.

Ogun, O.J., Thaller, G., Becker, D. (2022). An overview of the importance and value of porcine species in sialic acid research. Biology, 11(6), 903. DOI: 10.3390/biology11060903.

Ogun, O.J., Soremekun, O.S., Thaller, G., Becker, D. (2023a) An in silico functional analysis of non-synonymous SNPs of bovine CMAH gene and potential implication in pathogenesis. Pathogens, 12, 591. DOI: 10.3390/pathogens12040591.

Paysan-Lafosse, T., Blum, M., Chuguransky, S., Grego, T., Lázaro Pinto, B., Salazar, G.A., Bileschi, M.L., Bork, P., Bridge, A., Colwell, L., Gough, J., Haft, D.H., Letunić, I., Marchler-Bauer, A., Mi, H., Natale, D.A., Orengo, C.A., Pandurangan, A.P., Rivoire, C., Sigrist, C.J.A., Sillitoe, I., Thanki, N., Thomas, P.D., Tosatto, S.C.E., Wu, C.H., Bateman, A. (2023) InterPro in 2022. Nucleic Acids Research, 51, D418–D427. DOI: 10.1093/nar/gkac993.

Poulsen, N.A., Robinson, R.C., Barile, D., Larsen, L.B., Buitenhuis, B. (2019). A genome-wide association study reveals specific transferases as candidate loci for bovine milk oligosaccharides synthesis. BMC Genomics, 20, 404. DOI: 10.1186/s12864-019-5786-y.

Pupko, T., Bell, R.E., Mayrose, I., Glaser, F., Ben-Tal, N. (2002) Rate4Site: an algorithmic tool for the identification of functional regions in proteins by surface mapping of evolutionary determinants within their homologues. Bioinformatics, 18, S71–S77. DOI: 10.1093/bioinformatics/18.suppl_1.S71.

Samraj, A.N., Pearce, O.M.T., Läubli, H., Crittenden, A.N., Bergfeld, A.K., Banda, K., Gregg, C.J., Bingman, A.E., Secrest, P., Diaz, S.L., Varki, N.M., Varki, A. (2015). A red meat-derived glycan promotes inflammation and cancer progression. Proceedings of the National Academy of Sciences of the United States of America, 112(2), 542–547. DOI: 10.1073/pnas.1417508112.

Sato, C., Hane, M., Kitajima, K. (2016). Relationship between ST8SIA2, polysialic acid and its binding molecules, and psychiatric disorders. Biochimica et Biophysica Acta - General Subjects, 1860(8), 1739–1752. DOI: 10.1016/j.bbagen.2016.04.015.

Schauer, R. (2000). Achievements and challenges of sialic acid research. Glycoconjugate Journal, 17(7–9), 485–499. DOI: 10.1023/A:1011062223612.

Sharma, R., Ahlawat, S., Sharma, H., Aggarwal, R.A.K., Sharma, V., Tantia, M.S. (2019). Variable sialic acid content in milk of Indian cattle and buffalo across different stages of lactation. Journal of Dairy Research, 86(1), 98–101. DOI: 10.1017/S002202991800081X.

Sperandio, M., Frommhold, D., Babushkina, I., Ellies, L.G., Olson, T.S., Smith, M.L., Fritzsching, B., Pauly, E., Smith, D.F., Nobiling, R., Linderkamp, O., Marth, J.D., Ley, K. (2006). α2,3-Sialyltransferase-IV is essential for L-selectin ligand function in inflammation. European Journal of Immunology, 36(12), 3207– 3215. DOI: 10.1002/eji.200636157.

Tajima, F. (1989) Statistical method for testing the neutral mutation hypothesis by DNA polymorphism. Genetics, 123(3), 585–595. DOI: 10.1093/genetics/123.3.585.

Talenti, A., Powell, J., Hemmink, J.D., Cook, E.A.J., Wragg, D., Jayaraman, S., Paxton, E., Ezeasor, C., Obishakin, E.T., Agusi, E.R., Tijjani, A., Amanyire, W., Muhanguzi, D., Marshall, K., Fisch, A., Ferreira, B.R., Qasim, A., Chaudhry, U., Wiener, P., Toye, P., Morrison, L.J., Connelley, T., Prendergast, J.G.D. (2022). A cattle graph genome incorporating global breed diversity. Nature Communications, 13, 910. DOI: 10.1038/s41467-022-28605-0.

Tangvoranuntakul, P., Gagneux, P., Diaz, S., Bardor, M., Varki, N., Varki, A., Muchmore, E. (2003). Human uptake and incorporation of an immunogenic nonhuman dietary sialic acid. Proceedings of the National Academy of Sciences of the United States of America, 100(21), 12045–12050. DOI: 10.1073/pnas.2131556100.

UniProt Consortium (2023) UniProt: the Universal Protein Knowledgebase in 2023. Nucleic Acids Research, 51, D523–D531. DOI: 10.1093/nar/gkac1052.

Varki, N.M., Varki, A. (2007). Diversity in cell surface sialic acid presentations: Implications for biology and disease. Laboratory Investigation, 87, 851–857. DOI: 10.1038/labinvest.3700656.

Varki, A., Schauer, R. (2009). Sialic acids. In A. Varki, R.D. >Cummings, J.D. Esko, H.H. Freeze, P. Stanley, C.R. Bertozzi, G.W. Hart, M.E. Etzler (Eds.), Essentials of Glycobiology (2nd ed.). Cold Spring Harbor Laboratory Press.

Wang, B. (2012). Molecular mechanism underlying sialic acid as an essential nutrient for brain development and cognition. Advances in Nutrition, 3(3), 465S–472S. DOI: 10.3945/an.112.001875.

Weir, B.S., Cockerham, C.C. (1984) Estimating F-statistics for the analysis of population structure. Evolution, 38(6), 1358–1370. DOI: 10.2307/2408641.

Wickramasinghe, S., Hua, S., Rincon, G., Islas-Trejo, A., German, J.B., Lebrilla, C.B., Medrano, J.F. (2011) Transcriptome profiling of bovine milk oligosaccharide metabolism genes using RNA-sequencing. PLoS ONE, 6(4), e18895. DOI: 10.1371/journal.pone.0018895.

Wickramasinghe, S., Medrano, J.F. (2011) Primer on genes encoding enzymes in sialic acid metabolism in mammals. Biochimie, 93(10), 1641–1646. DOI: 10.1016/j.biochi.2011.06.002.

